# Joint Segmental Duplication Co-option Drives Human-specific Transcriptional Readthrough and Expression Fine-tuning of *NPEPPS*-*TBC1D3*

**DOI:** 10.64898/2026.01.14.699191

**Authors:** Kaiyue Ma, Zikun Yang, Zhengtong Li, Jingyi Guo, Da Lian, Ziyi Wang, Hongyi Ma, Shilong Zhang, Lianting Fu, Hezheng Lyu, Xinrui Jiang, Qiujin Xie, Guanting Li, Chenhao Yang, Jieyi Chen, Juan Zhang, Peng Liu, Xiangyu Yang, Zhen-Ge Luo, Guojie Zhang, Yafei Mao

**Affiliations:** Bio-X Institutes, Key Laboratory for the Genetics of Developmental and Neuropsychiatric Disorders, Ministry of Education, International Peace Maternity and Child Health Hospital, Shanghai Key Laboratory of Embryo Original Diseases, School of Medicine, Shanghai Jiao Tong University, Shanghai, China; Zhiyuan College, Shanghai Jiao Tong University, Shanghai, China; Division of Chemistry I, Department of Medical Biochemistry and Biophysics, Karolinska Institutet, Stockholm, Sweden; Veterinary Hospital, Shanghai Wild Animal Park, Shanghai, China; School of Life Science and Technology & State Key Laboratory of Advanced Medical Materials and Devices, ShanghaiTech University, Shanghai, China; Center for Genomic Research, International Institutes of Medicine, Fourth Affiliated Hospital, Zhejiang University, Yiwu, China; Center of Evolutionary & Organismal Biology, and Women’s Hospital at Zhejiang University School of Medicine, Zhejiang University, Hangzhou, China; School of Medicine, Zhejiang University, Hangzhou, China; Center for Comparative Biomedicine, Ministry of Education Key Laboratory of Systems Biomedicine, State Key Laboratory of Medical Genomics, Institute of Systems Biomedicine, Shanghai Jiao Tong University, Shanghai, China

## Abstract

Segmental duplication (SD) is a major driver of functional changes in evolution and disease. Many genes embedded within SDs, such as *NPEPPS* and *TBC1D3*, display substantial copy number variation (CNV) across individuals. Yet, the precise identification of the functional copies and their transcriptional outputs remains largely unstudied. Focusing on *NPEPPS* and *TBC1D3*, we illustrate human-specific expression fine-tuning mechanisms associated with readthrough transcripts. We identified a human-specific *NPEPPS*-*TBC1D3* digenic genomic structure that originated from a joint SD pair and became fixed across populations. Experiments demonstrate that this structure generates *NPEPPS*-*TBC1D3* readthrough transcripts, which are the predominant isoforms of *TBC1D3* expression in various cell types, fine-tuning its protein level. Furthermore, a human-specific hypomethylation signal within an upstream CpG island of *NPEPPS* precisely pinpoints the expressed *TBC1D3* paralog. Moreover, we reveal transcriptional readthrough events are ∼3-fold enriched for joint-SD-associated transcriptional readthrough (JSDTR) and identify 109 JSDTR gene pairs, including neurodevelopmentally important pairs and clinically interesting *SERF1A/B-SMN1/2*. Taken together, our findings comprehensively describe an example of how a joint SD event shaped evolution and suggest that JSDTR is a broad mechanism for the emergence of new functions.

## Main

Segmental duplications (SDs) are large, low-copy repeats typically >1 kb in length and sharing >90% sequence identity^1^, which frequently incorporate into genomic regions through non-allelic homologous recombination (NAHR)^2^. The expansion of SDs can potentially create copy number variations (CNVs), new gene bodies, new transcripts (including read-through transcripts), and fused open reading frames (ORFs)^3,4^. While such intense reorganization of genomic elements has been recognized as crucial for forming human-specific genomic features important to evolution, previous studies focused primarily on human-specific genes^3,5,6^, failing to highlight the significant roles of human-specific transcriptional/translational regulatory mechanisms resulting from SD expansion.

Here, we report a joint SD locus in the human genome (chr17:39044723-39120263 on the T2T-CHM13 reference genome, “the 39M locus”), where two different SDs converge, bring *NPEPPS* and *TBC1D3* gene bodies into close proximity, and give rise to *NPEPPS-TBC1D3* readthrough transcripts. While *NPEPPS-TBC1D3* transcripts have been detected in previous studies^7,8^, their presence has been mistakenly attributed to disease-related genomic events^9^. In this study, we establish that *NPEPPS-TBC1D3* transcripts in fact originate from a joint SD locus fixed across human populations and are commonly expressed in various cell types of healthy human individuals as the predominant *TBC1D3* transcripts, representing a unique evolutionary case of gene regulation.

First, using genome assemblies from Human Pangenome Reference Consortium (HPRC) Phase 1^10^, Human Genome Structural Variation Consortium (HGSVC) Phase 3^11^, and Asian Pan-Genome Project (APG) Phase 1 (Wu *et al.*, manuscript in revision), as well as the T2T-CHM13^12^ and the CN1 diploid genome^13^, we have completely charted *NPEPPS* and *TBC1D3* paralogs across human populations, respectively (Fig. 1 and Supplementary Fig. 1,2). 462 haplotypes with a single contig covering all *TBC1D3* and *NPEPPS* loci were selected for subsequent analyses, avoiding contig breakpoints that can complicate the results.

**Fig. 1:**
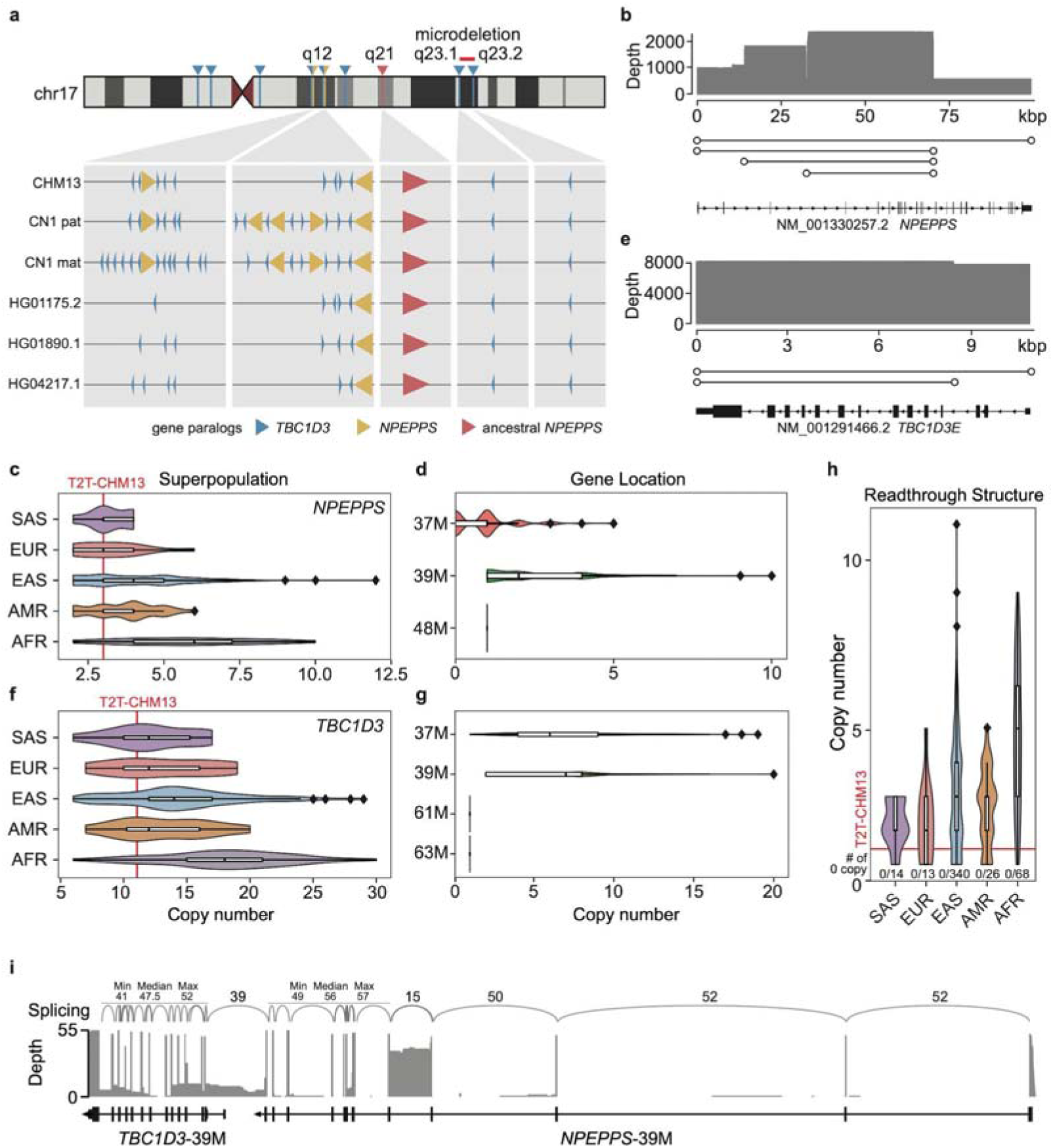
The joint SD-derived *NPEPPS-TBC1D3* digenic genomic structure is fixed across populations and generates *NPEPPS-TBC1D3* readthrough transcripts. **a**, Ideogram illustrating the genomic locations of *TBC1D3* and *NPEPPS* genes on human chromosome 17. Red, yellow and blue triangles represent ancestral *NPEPPS*, derived *NPEPPS* and *TBC1D3*, respectively. **b**, **c**, **d** Analysis of *NPEPPS* segmental duplication breakpoints (b) and copy number distribution grouped by superpopulation (c) and genomic location (d). **e**, **f**, **g** Analysis of *TBC1D3* segmental duplication breakpoints (e) and copy number distribution grouped by superpopulation (f) and genomic location (g). **h**, Copy number variation of the digenic structure across human populations shows at least one digenic structure per haplotype. **i**, Readthrough splicing profile from CHM13hTERT Iso-seq data. Curves represent splicing junctions, with numbers above indicating the number of supporting reads. Population abbreviations: SAS, Southern Asian (n=14 haplotypes). EUR, European (n=13 haplotypes). EAS, Eastern Asian (n=340 haplotypes). AMR, American (n=26 haplotypes). AFR, African (n=68 haplotypes).

NPEPPS has been reported to protect against TAU-induced neurodegenerative disorders, such as Alzheimer disease and amyotrophic lateral sclerosis, by proteolysis of TAU protein^14–16^. The orthologous ∼99.4-kbp *NPEPPS* gene body is located at human 17q21.32 (chr17:48385617-48485055, T2T-CHM13, “the 48M locus”). A previous study reported that incomplete segmental duplication of *NPEPPS* resulted in multiple ∼63.0-kbp paralogs^17^. In our study, we increased the completeness of the analysis, identifying various incomplete duplicates with different lengths across human populations (Fig 1a,b). Aside from the ∼100-kbp complete *NPEPPS* copy, incomplete copies exhibit lengths clustered around three peaks (∼70.2 kbp, 56.1 kbp and 37.3 kbp; determined by aligning the ∼100-kbp sequence to other loci). The 70.2-kbp copies have a complete 5’ end but missing the 3’ end (*e.g.*, RefSeq *NPEPPSP1* at the 39M locus; chr17:39058811-39120263), while other copies are truncated at both the 5’ and the 3’ ends (*e.g.*, RefSeq *LOC101060212* at the 37M locus; chr17:37222028-37267942) (Fig. 1a-d and Supplementary Fig. 3a). Together, *NPEPPS* exhibits high CNV in the human population. Here, we identified 2 to 12 *NPEPPS* copies per haplotype (Fig. 1c,d and Supplementary Table 1A), corresponding to the previously reported range of 4-12 copies per diplotype^18^.

*TBC1D3*, on the other hand, encodes TBC1 domain family member 3, which, while reported to be involved in Rab GTPase signaling, has its detailed roles and associated pathways under debate^19,20^. Despite many exciting functions proposed and reported^21–23^, significant gaps remain in our understanding of the regulation of *TBC1D3* expression, casting doubts on the biological relevance of previous functional results. The *TBC1D3* gene family is characterized by a complex pattern of SDs, resulting in two distinct groups of paralogs. One includes the dispersed paralogs that were annotated as pseudogenes (*TBC1D3P1* to *P5* and *P7*, at chr17 61M, 63M, 20M, 18M, 28M, and 42M; *TBC1D3P6*, at chr1 15M). In contrast, the other group comprises the complete and potentially functional copies that are tightly clustered in the 37M and 39M loci (Fig. 1a)^24^. Here, we identified 6-30 copies per haplotype (Fig. 1e-g and Supplementary Table 1A), corresponding to the previously reported range of 23-42 per diplotype^18^.

In contrast to *NPEPPS*, most duplicates of *TBC1D3* (95.1%, 6523/6858) retain the 10.9-kbp complete gene body (Fig. 1e and Supplementary Fig. 3b), while a small fraction (4.8%, 331/6858) possesses a 8.4-kbp gene body that lacks the first three exons (Supplementary Table1B,C). Despite the extensive CNV, human *TBC1D3* expression was reported to be primarily contributed by a single paralog group: *TBC1D3-CDKL*^24^, one of the two expanded *TBC1D3* clusters at 17q12 (Fig. 1a). Although this observation that only one cluster is the main contributor of *TBC1D3* expression is biologically reproducible, the accurate attribution to a specific *TBC1D3* paralog was previously not achieved. Instead, previous attributions are largely influenced by different gene annotation processes, lacking a sequence- and context-aware analysis (Supplementary Fig. 4). Therefore, we sought to pinpoint the exact paralog(s) that drives the transcription and subsequent translation of TBC1D3.

The substantial variations of *NPEPPS* and *TBC1D3* motivated us to examine the joint SDs of these two genes and their effects on the regulation of TBC1D3. We first examined the 48M locus where the canonical *NPEPPS* resides and detected no joint SD. The *NPEPPS* at this locus is with fixed orientation and CN of 1 (462/462 haplotypes) (Fig. 1a,d and Supplementary Fig. 1,2), and no *TBC1D3* paralog is found near *NPEPPS*-48M (± 1Mbp) in all assemblies. Similarly, no *NPEPPS* paralog is found near any of *TBC1D3P1* to *TBC1D3P7* (± 1Mbp), correlating with our Iso-Seq results, which show no evidence of expression for these paralogs (Supplementary Fig. 5a,b). Interestingly, a previous study reported a recurrent ∼2.2-Mbp microdeletion at 17q23.1-q23.2 flanked by a pair of SDs<u>(Ballif et al., 2010)</u>. The deletion of *TBX2* and *TBX4* results in heart defects and limb abnormalities. We identified *TBC1D3P1* and *TBC1D3P2* as the SDs flanking this region, revealing their role as the structural basis for this recurrent microdeletion (Supplementary Fig. 5c-e).

In contrast to the loci examined above, the 37M and the 39M loci contain both *NPEPPS* (37M: 0-5 copies per haplotype; 39M: 1-10 copies) and *TBC1D3* paralogs (37M: 1-19 copies; 39M: 2-20 copies). While the orientation and CN of the *NPEPPS* paralogs vary at the 39M locus, all assemblies possess at least one *NPEPPS* paralog (hereafter, “*NPEPPS*-39M”) (Fig. 1a,c,d,f,g, Supplementary Fig. 1,2, and Supplementary Table 1B). Most importantly, in all haplotypes, a *TBC1D3* paralog with the same orientation as *NPEPPS*-39M can be found at its immediate downstream region, indicating that the *NPEPPS*-*TBC1D3* digenic structure originated from joint SDs and became fixed across human populations (Fig. 1a,h, Supplementary Fig. 1,2, and Supplementary Table 1B). The similar digenic structure also exists for a subset of other paralogs at the 39M and the 37M loci, indicating duplications of the joint SD sequence as a whole unit following the convergence of the respective SDs (Fig. 1a,h and Supplementary Fig. 1,2).

Importantly, Iso-Seq data from the CHM13hTERT cell line reveals expression of *NPEPPS*-*TBC1D3* readthrough transcripts at places with the digenic structure (Fig. 1i). In the readthrough transcripts, the terminal exon of *NPEPPS*-39M is partially skipped and spliced to the second exon of *TBC1D3*-39M, preserving the canonical start codon for coding TBC1D3. This splice junction is supported by 39 reads, comparable to the support read counts observed for other junctions within *NPEPPS*-39M and *TBC1D3*-39M (min=15, median=50, max =57 reads, n = 23) (Fig. 1i and Supplementary Table 2A). Furthermore, we observed no read supporting the canonical splicing event between the first and second exons of *TBC1D3*-39M. This absence likely indicates that the independent transcription of *TBC1D3*-39M occurs at such a low proportion that it falls below the detection limit of the Iso-Seq experiments. To validate the existence of this readthrough splicing in natural human cellular contexts, we generated induced pluripotent stem cells (iPSCs), iPSC-derived neural progenitor cells (NPCs), and iPSC-derived forebrain neurons (FBNs) using the blood sample from a healthy donor (CN1) (Supplementary Fig. 6a). *NPEPPS*-*TBC1D3* readthrough transcripts are identified in all three cell types (Supplementary Fig. 6b). Taken together, the evidence illustrates that *NPEPPS*-*TBC1D3* readthrough transcripts naturally occur across diverse cell types in healthy individuals.

Next, we establish *NPEPPS*-*TBC1D3* are the predominant transcripts responsible for *TBC1D3* expression in human neural lineage cells, and potentially other cell types. A *TBC1D3*-specific primer (*TBC1D3*-Rev1) was designed and utilized in a 5’ Rapid Amplification of cDNA Ends (5’ RACE) experiment (Fig. 2a, see also Supplementary Fig. 7a,b for the pilot experiment using *TBC1D3*-Rev0). TOPO cloning using the 5’ RACE product exhibits *NPEPPS*-*TBC1D3* as the predominant transcripts of *TBC1D3* in NPCs (18 of 20 colonies; 1 non-specific; 1 noise; 0 non-readthrough) (Supplementary Fig. 7c,d). Interestingly, the *NPEPPS* part of these readthrough transcripts exhibits high diversity, with individual transcripts containing different ORFs. This variation likely results from variable splicing events (Supplementary Fig. 7e, 8-11).

**Fig. 2:**
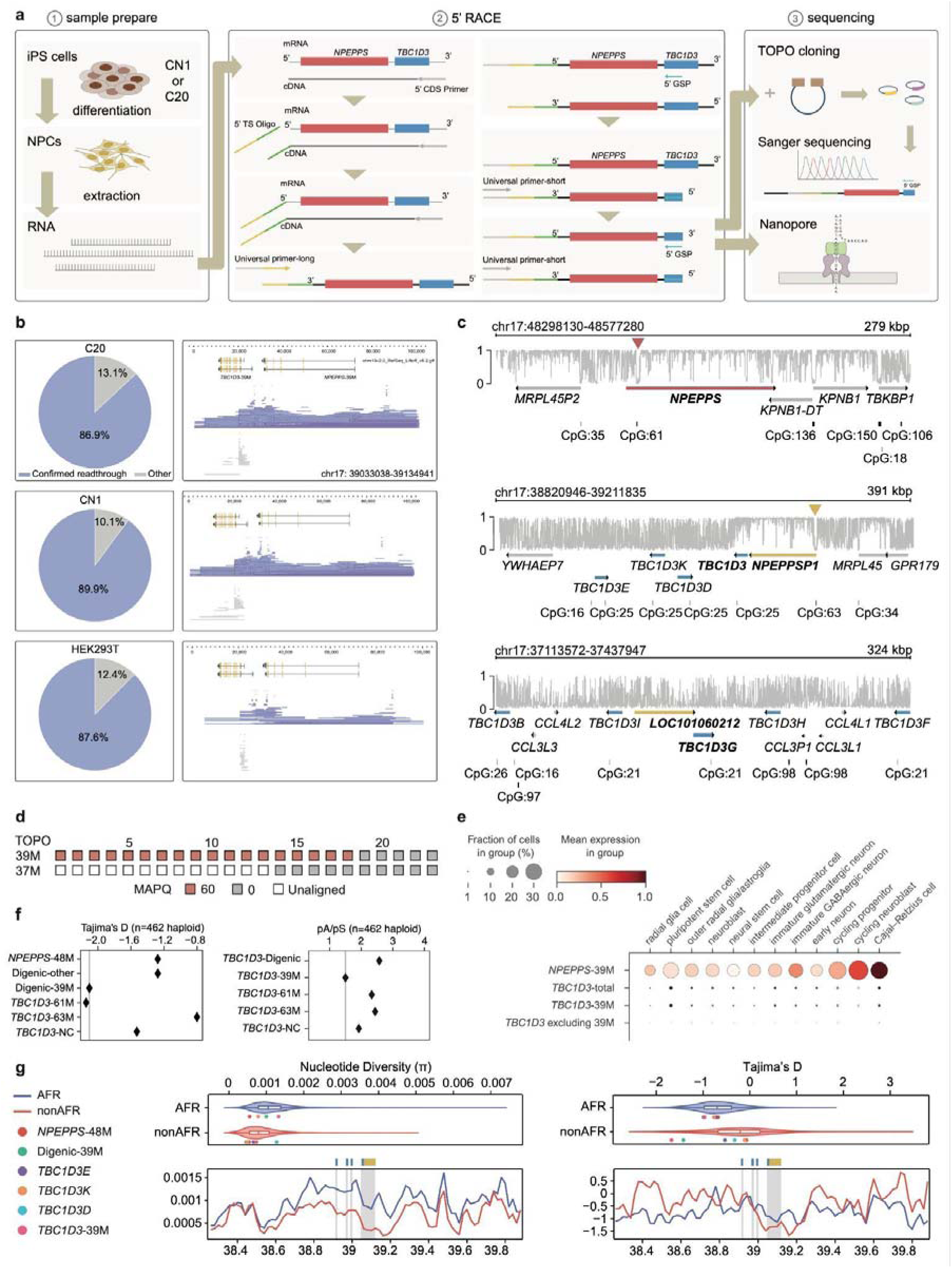
The *NPEPPS-TBC1D3* transcripts, specifically those from the 39M locus, are the predominant source of *TBC1D3* expression in various cell types. **a**, RNA was extracted from CN1 NPC and C20 NPC, and 5’ RACE was performed to determine the transcription starting sites of the transcripts using the *TBC1D3* Gene Specific Primer. 5’ RACE products were evaluated with both TOPO-Sanger sequencing or nanopore sequencing. **b**, Proportions of confirmed readthrough and other reads detected by 5’ RACE-nanopore in C20 NPC, CN1 NPC, and HEK293T cells, along with JBrowse2 read cloud displays showing split-read support at the *NPEPPS*-*TBC1D3* junction, confirming the predominance of readthrough transcripts. **c**, The CpG island annotation and methylation levels of the 48M, 39M and 37M loci. At the 48M and the 39M loci, the hypomethylated CpG island resides at the upstream of *NPEPPS*. In contrast, the 37M locus does not harbor such a CpG island. **d**, The mapping quality (MAPQ) of 23 reconstructed transcript sequences from TOPO-cloning demonstrates that readthrough transcript sequences are more similar to the 39M sequence than the 37M sequence, indicating the readthrough expression is mainly contributed by the 39M locus. **e**, Single-cell analysis of human brain organoid samples (CN1 & C50, Day 42 & Day 98, combined) shows *TBC1D3*-39M is the predominant paralog for *TBC1D3* expression. **f**, The Tajima’s D and pA/pS of *NPEPPS* and *TBC1D3* duplicates indicates that *TBC1D3*-39M is under selection on both nucleotide and amino acid level in the human population (n=462 haploids). Digenic, the *NPEPPS*-*TBC1D3* digenic region. NC, negative control, representing the *TBC1D3* copies excluding *TBC1D3*-39M. **g**, Nucleotide diversity (π) and Tajima’s D analysis of *TBC1D3*-39M based on HGSVC3-called single nucleotide variants (n=65). No significant reduction in nucleotide diversity was observed in African populations (p=0.439) or non-African populations (p=0.685). Similarly, no significant deviation in Tajima’s D values was detected in African populations (p=0.204) or non-African populations (p=0.113), when compared to genome-wide averages calculated from 80 kbp sliding windows across chromosome 17. This result, however, is more likely a product of methodological limitations than an actual evolutionary trend.

To accurately quantify the proportion of *NPEPPS*-*TBC1D3* in the 5’ RACE product, nanopore sequencing was performed, which again confirms the majority of the *TBC1D3* transcripts are *NPEPPS*-*TBC1D3* transcripts (>86.9% in C20 NPC; >89.9% in CN1 NPC; >87.6% in HEK293T) (Fig. 2b, Supplementary Fig. 12, and Supplementary Table 2B). This conclusion is supported by estimation using short-read RNA-seq data from CN1 iPSC (∼91.1%-91.4%), NPC (∼67.6%-91.9%), and FBN (∼86.0%-98.2 %) (Supplementary Fig. 6b, 13a). Interestingly, testis is another tissue with reported high *TBC1D3* expression. Using publicly available RNA-seq data (PRJNA1092349), we estimate that the readthrough transcripts contribute to ∼29.3%-56.6% of the *TBC1D3* expression in healthy testicular tissues, which potentially suggests different regulation mechanisms from the neurodevelopmental regulation (Supplementary Fig. 13).

Given that the digenic structure is present at the 39M locus in all individuals but absent at the 37M locus in a subset of individuals, we hypothesize that the 39M locus is the primary locus for *NPEPPS*-*TBC1D3* expression. To validate this hypothesis, we performed a methylation analysis, which reveals a hypomethylation signal in a CpG island upstream of *NPEPPS*-39M (chr17:39119773-39120426). A similar signal also presents upstream of *NPEPPS*-48M (chr17:48392644-48393344), but not *NPEPPS*-37M due to the lack of the CpG island sequence (Fig. 2c and Supplementary Fig. 14). Additionally, the readthrough transcript sequences (reconstructed from TOPO-Sanger sequencing) are more similar to *NPEPPS*-39M than *NPEPPS*-37M based on the mapping quality (MAPQ) scores. Among 23 transcript sequences, eighteen have higher MAPQ for *NPEPPS*-39M while the remaining five sequences exhibited equal mapping confidence to both loci (MAPQ = 0), indicating ambiguous alignment (Fig. 2d and Supplementary Table 2C). Furthermore, single-cell analysis of human brain organoid samples at Day 42 and Day 98 shows again *TBC1D3*-39M is the predominant paralog for *TBC1D3* expression (Fig. 2e and Supplementary Fig. 15).

To examine the constraint of this loci while avoiding introducing false negative signals from other copies, we analyzed the Tajima’s D and pA/pS of the ancestral *NPEPPS*, *NPEPPS*-39M, *TBC1D3*-39M, *TBC1D3*-61M, *TBC1D3*-63M and other copies from each haplotype. Our results show that the 39M *NPEPPS*-*TBC1D3* digenic region and *TBC1D3*-61M have significant negative Tajima’s D and may experience recent selective sweeps. Moreover, while all pA/pS values are beyond 1 (meaning positive selection), *TBC1D3*-39M shows the lowest value among all *TBC1D3* copies, indicating this locus is more conserved on amino acid level compared with other copies. In general, our constraint analyses indicate that *TBC1D3*-39M is the most conserved paralog at both nucleotide and amino acid levels, supporting the notion that it is the main functional paralog in the *TBC1D3* family (Fig. 2f). In addition to our approach, we explored employing previously commonly-used methods to compute nucleotide diversity (π) and Tajima’s D on the whole chromosome level, using long-read-based variants from HGSVC3. However, nucleotide diversity (π) doesn’t exhibit significantly lower values in both African (p=0.439) and non-African populations (p=0.685) (Fig. 2g). Similarly, Tajima’s D is also not significantly lower in African (p=0.204) and non-African populations (p=0.113) (Fig. 2g). We speculate that this result is probably due to limitations of these methods, specifically, the ambiguous alignment of reads from different loci when dealing with duplicated genes, which introduce false negative results. Collectively, the observations above support that the cis-regulatory elements of *NPEPPS*-39M are major drivers of the expression of the *NPEPPS*-*TBC1D3* transcripts.

Notably, while *TBC1D3* has been frequently referred to as a hominoid-specific gene^19,23^, its gene sequence is in fact commonly present in other simians^24–26^. Similar as in humans, the gene expression and regulation of *TBC1D3* across simian species remain largely understudied. Using primate telomere-to-telomere (T2T) genomes^27,28^, we have elucidated the evolutionary history of *NPEPPS* and *TBC1D3*, shedding light on the evolutionary path of the *NPEPPS*-*TBC1D3* digenic region (Fig. 3a). Macaques and orangutans have one single *NPEPPS* copy (syntenic to the 48M locus) per haploid genome (Supplementary Table 3A,B). This ancestral *NPEPPS* is located adjacent to an inversion–translocation breakpoint that distinguishes orangutans from hominines (African great apes) (Fig. 3a). These structural rearrangements potentially led to the partial duplication resulting in the additional gene copy similar to *NPEPPS*-39M (Fig. 3b).

**Fig. 3:**
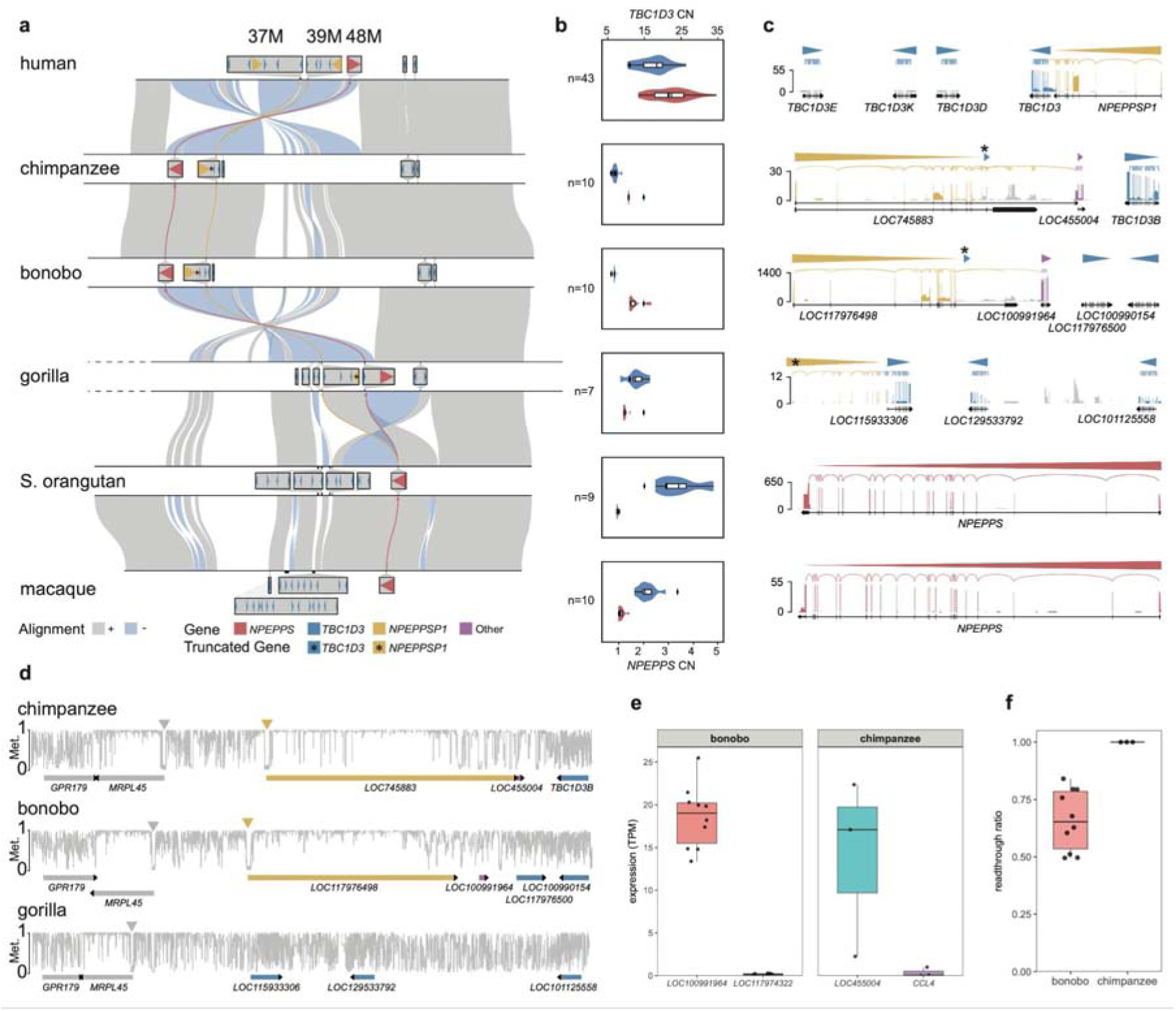
Evolutionary analyses reveal that the intact *NPEPPS*-*TBC1D3* sequence is human-specific and the *NPEPPS* readthrough occurs with different genes in different primates. **a,** Synteny comparison of human chromosome 17 and its NHP counterparts illustrates the evolutionary trajectory of *NPEPPS* and *TBC1D3*. **b,** Copy number estimation based on FastCN shows the expansion of *NPEPPS* copies in African great apes and expansions of *TBC1D3* copies in different lineages. **c,** Iso-Seq reveals expression of readthrough transcripts, including *NPEPPS*-*TBC1D3* in humans and *NPEPPS*-*CCL4* in bonobos and chimpanzees. **d,** DNA CpG methylation analysis indicates that hypomethylated CpG islands exist in the upstream of the *NPEPPS*, which is syntenic to human *NPEPPS*-39M, in chimpanzees and bonobos. The *NPEPPS* sequence in this syntenic region in gorillas is substantially truncated and not annotated as a gene by RefSeq. Consequently, no such hypomethylation signal is observed in gorillas. Met., Methylation level. **e**, Expression levels of duplicated *CCL4* gene copies in bonobo and chimpanzee. In each species, one copy forms readthrough transcripts with the upstream *NPEPPS* gene (*NPEPPS-LOC100991964* in bonobos and *NPEPPS-LOC455004* in chimpanzees), whereas the other copy does not. Individual dots represent independent bulk RNA-seq samples. The bonobo data were obtained from publicly available prefrontal cortex bulk RNA-seq datasets (PRJEB33938). The chimpanzee data were generated in this study using NPCs, including two independent differentiation batches from the individual SY2-Z2 and one batch from the individual SY2-D2. In both species, the *NPEPPS-CCL4* readthrough-associated copy shows higher expression compared with the other *CCL4* copy. **f**, Estimated proportion of *NPEPPS-CCL4* readthrough transcripts relative to all *CCL4* transcripts at the readthrough loci in bonobos and chimpanzees. The readthrough ratio is estimated as the proportion of readthrough-supporting split-read depth relative to the total read depth at a representative genomic position. Each dot represents an independent bulk RNA-seq sample.

Importantly, while the SDs of *TBC1D3* and *NPEPPS* are spatially associated in all hominines, the *NPEPPS*-*TBC1D3* digenic structure, where those two genes are immediately neighboring and share the same orientation, is fully intact only in humans (Fig. 3c). In chimpanzees and bonobos, the downstream *TBC1D3* is heavily truncated and consequently precludes the *NPEPPS*-*TBC1D3* readthrough transcripts. Interestingly, the partially duplicated *NPEPPS* generates readthrough transcripts together with its more distal gene: *NPEPPES*-*LOC455004* in chimpanzee and *NPEPPES*-*LOC100991964* in bonobo. Both partner genes are annotated as immune-related C-C motif chemokine 4 (*CCL4*) genes (Fig. 3c,d and Supplementary Fig. 16). Moreover, both chimpanzees and bonobos have two copies of *CCL4*, and in both species, the copy that forms the *NPEPPS*-*CCL4* readthrough transcript has higher expression compared to the other in neural lineage cells (Fig. 3e and Supplementary Table 3C). At the *NPEPPS*-*CCL4* locus, the readthrough transcripts are estimated to constitute ∼49.5% to ∼100% of all *CCL4* transcripts (Fig. 3f). On a separate note, in gorillas, while *TBC1D3* retains the intact gene structure, the upstream *NPEPPS* copy is truncated at its 5’ end, thereby lacking the CpG island sequence seen in human *NPEPPS*-39M (Fig. 2c, 3c, 3d, and Supplementary Fig. 16). Taken together, the intact *NPEPPS*-*TBC1D3* digenic structure, with the complete *TBC1D3* ORF sequence and the hypomethylated *NPEPPS* CpG island, is specific to humans. This unique genomic configuration may underlie human-specific expression patterns of *TBC1D3* (Supplementary Fig. 17).

Furthermore, we hypothesize that the *NPEPPS-TBC1D3* digenic structure serves to fine-tune the TBC1D3 protein abundance through post-transcriptional and translational mechanisms, controlling the TBC1D3 level to the physiological optimal. To explore this hypothesis, we cloned several different (*NPEPPS*-) *TBC1D3* transcripts (Fig. 4a and Supplementary Fig. 18). Similar to what is observed in the 5’ RACE analysis, the transcripts exhibit substantial splicing variability, both in the *NPEPPS* region and in the *TBC1D3* region, adding another layer of complexity to the post-transcriptional and translational regulation. Using a pCAGGS expression system in HEK293T cells, plasmids constructed using cDNA of the readthrough transcripts generate lower TBC1D3 protein level compared to what is generated by *TBC1D3*-only cDNA sequences (Fig. 4b,c and Supplementary Fig. 19a,b).

**Fig. 4:**
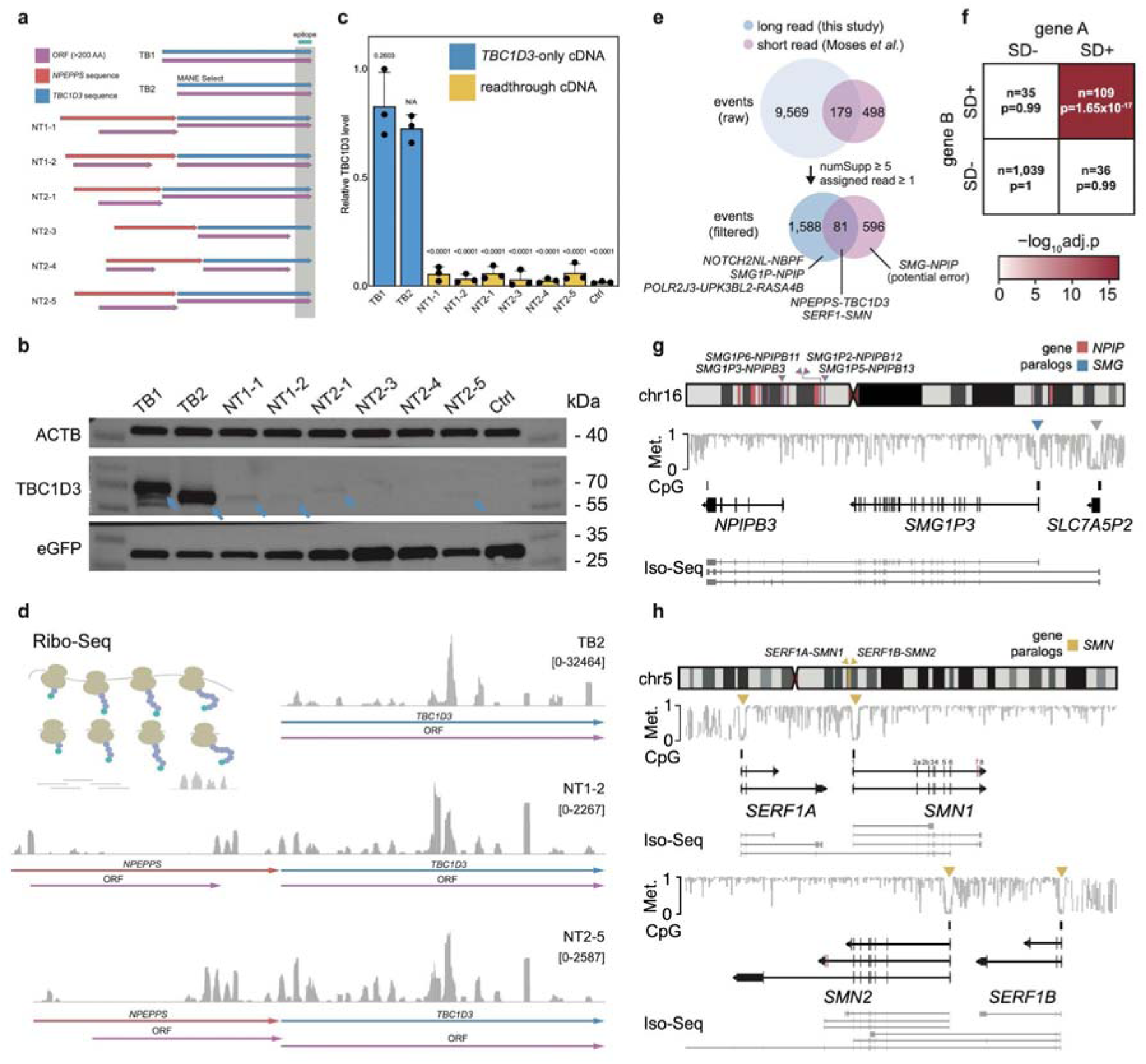
The *NPEPPS-TBC1D3* transcripts fine-tune the TBC1D3 protein level, and joint-SD-related readthrough events broadly exist. **a**, (*NPEPPS*-) *TBC1D3* transcripts were cloned into a pCAGGS expression system. The position of the epitope of the anti-TBC1D3 antibody is indicated. TB2 represents the MANE Select isoform. **b**,**c**, The readthrough transcripts generate lower TBC1D3 protein level compared to *TBC1D3*-only sequences. NT2-3 and NT2-4 are expected to generate no band because of the lack of the epitope sequence. Blue arrows: TBC1D3 bands. ANOVA and Dunnett’s multiple comparisons tests were performed for TB2 *vs*. each of the other samples. **d**, Ribo-Seq was performed for TB2, NT1-2, and NT2-5. Regularly distributed peaks were observed in the whole span of the *TBC1D3* ORF but not the whole span of the *NPEPPS* ORFs. **e**, Benchmark between our long-read-based fusion calling results and the short-read-based results from Moses *et al.*, 2025 shows that *NPEPPS*-*TBC1D3* is a commonly identified gene fusion pair. **f**, Enrichment analysis between SD-associated gene pairs and fusion gene pairs using Fisher’s exact test and Benjamini-Hochberg method shows that transcriptional readthrough events exhibit significant enrichment of JSDTR gene pairs (p-value=1.65e-17). **g**, Diagram of *SMG1P3*-*NPIPB3* localization, methylation and representative reads on chr16. **h**, Diagram of *SERF1A*/*B*-*SMN1*/*2* localization, methylation and representative reads. Among all readthrough reads, 30.2% (13/43) and 60.5% (26/43) terminate respectively a few bases after exon 3 (chr5:71,399,360 for *SMN1*; chr5:70,820,191 for *SMN2*) and after exon 6 (chr5:71,403,067 for *SMN1*; chr5:70,816,484 for *SMN2*). 6.98% (3/43) and 2.32% (1/43) readthrough reads terminate at exon 8 with exon 7 inclusion and skipping, respectively.

It is likely because the *NPEPPS* sequence within a readthrough transcript can be deemed an ultra-long 5’ UTR of *TBC1D3* (Supplementary Fig. 19c-f and Supplementary Table 2D), notably longer than what is typically observed in mRNAs^29,30^, thereby adding the chance of decreased cap-dependent translation efficiency. For instance, many *NPEPPS*-*TBC1D3* transcripts are polycistronic, containing ORFs within the *NPEPPS* sequence upstream of the *TBC1D3* sequence (Fig. 4 and Supplementary Fig. 8-11, 18), which may further lower the translation efficiency^31–33^. It is worth noting that an internal ribosome entry site (IRES) sequence, if exists, could enhance the cap-independent translation efficiency of the *TBC1D3* sequence within the readthrough transcripts; however, we were unable to detect such a sequence (Supplementary Fig. 20). Furthermore, the versatile splicing schemes potentially introduce an additional layer of control to the 5’ UTR of these transcripts (Supplementary Fig. 7-11, 18). Additionally, we cannot rule out inhibitory transcriptional regulatory effects, as the RNA level of the readthrough cDNA is also lower than that of the *TBC1D3*-only cDNA (Supplementary Fig. 21).

Next, to better examine the translation of the readthrough transcripts, using the same pCAGGS system, we performed Ribo-Seq for two representative readthrough transcripts (NT1-2 and NT2-5) and a *TBC1D3*-only transcript. While the *TBC1D3* ORF in the readthrough transcripts exhibits regularly distributed peaks, sharing the same pattern with the ORF in the *TBC1D3*-only cDNA, Ribo-Seq peaks within the *NPEPPS* region do not cover the entire ORFs, suggesting the selected representative readthrough transcripts mainly execute TBC1D3 functions (Fig. 4d and Supplementary Fig. 22).

Finally, using Iso-Seq data from CHM13hTERT, CN1 NPC, and C20 NPC, we identified 9,569 readthrough events with at least one Full-Length Non-Chimeric (FLNC) read supported. After quality control (Methods), we retained 1,588 high-confidence fusion events and 1,219 unique gene pairs (Fig. 4e and Supplementary Table 4A). Among them, 109 are associated with joint-SD-associated transcriptional readthrough (JSDTR). Transcriptional readthrough events exhibit significant enrichment of JSDTR gene pairs (p-value=1.65e-17), but not other pairs (p-values > 0.99), with an average 2.63-fold difference (Fig. 4f and Supplementary Table 4B). Among the JSDTR gene pairs, we identified previously detected *NOTCH2NLA*/*B*/*C*-*NBPF10*/*14*/*19* and *ARHGAP11B*-*LOC100288637*^34^ (Supplementary Fig. 23). Similar to *NPEPPS*-*TBC1D3*, the downstream partners, *NBPF*s and *LOC100288637,* lack CpG islands at 5’ ends of their gene models, whereas the upstream partners, *NOTCH2NL*s and *ARHGAP11B*, have hypomethylated CpG islands in their promoter regions. Interestingly, the RefSeq gene model of *NBPF26* (chr1:120,737,164-120,851,627) contains *NOTCH2NLR* (chr1:120,737,164-120,808,094) within its 5’ end, reflecting, to some extent, the process of fusion gene birth. Additionally, a recent study highlighted the neurodevelopmental function of the gene pair of *NOTCH2NLB* and *NBPF14*, where they were described as co-evolving and co-expressing^35^. We speculate the *NOTCH2NLB*-*NBPF14* readthrough transcripts may also contribute to neurodevelopmental functions, pending further experimental validation.

Importantly, here, we also identified other biomedically relevant gene pairs that have previously not been highlighted, including *SMG1P2*/*P3*/*P5*/*P6*-*NPIPB12*/*B3*/*B13*/*B11* and *SERF1A*/*B*-*SMN1*/*2* (Fig. 4g,h and Supplementary Fig. 23). Previously, the role of *NPIP* paralogs in genomic rearrangements and their fusion with *PKD1* paralogs (*PKD1*-*NPIPA*) were reported^36^. A recent study using GRCh38 detected the fusion of *SMG1* and *NPIPB5*^37^; however, these 2 genes have opposite orientations and are separated by approximately 3.6 Mbp in hg38. While this “*SMG1*-*NPIPB5*” fusion was reported to be universal across hundreds of individuals, this observation likely arises from mis-mapping between highly similar paralogous copies using short-read RNA-seq data and fails to be validated using Iso-Seq data. In contrast, our Iso-Seq-based analysis identified four fusion pairs involving *SMG1* pseudogenes and *NPIP* paralogs (average gene pair distance: ∼ 6 kbp) (Fig. 4g and Supplementary Fig. 23). Again, the downstream *NPIP* partners lack CpG islands near the 5’ ends of their gene models, while their upstream *SMG1* partners contain hypomethylated CpG islands except for *SMG1P6*. In the case of *SMG1P6*, a hypomethylated CpG island can be found in the promoter region of its immediate upstream gene *BOLA2*, generating *BOLA2*-*SMG1P6*-*NPIPB11* transcripts. These examples collectively demonstrate that the co-option of regulatory elements of neighboring genes is a key mechanism by which duplicated genes achieve expression.

Additionally, JSDTR gene pairs, such as *SERF1B*-*SMN2*, are clinically relevant (Fig. 4h). *SMN1* and *SMN2* are associated with Spinal Muscular Atrophy (SMA), which is caused by abnormality of *SMN1* and modified by the varying copy numbers of *SMN2*^38^. It is biomedically interesting to study whether *SERF1*-*SMN* readthrough transcripts modify disease phenotypes and thereby have implications for diagnostics and treatment development. On the one hand, while a previous study has investigated the modifying effects of *SERF1A*^39^, the analyses remain to be refined by accounting for the *SERF1*-*SMN* readthrough transcripts. On the other hand, one of the main disease-modifying treatments employs antisense oligonucleotides or small molecules to promote *SMN2* exon 7 inclusion to restore the level of full-length functional SMN protein^40^. Future research is needed to determine how *SERF1B*-*SMN2* reacts to these treatments. Notably, unlike the examples showcased above, hypomethylated CpG islands can be found in the promoter regions of both *SERF1A*/*B* and *SMN1*/*2*, indicating a more complex regulatory system.

In summary, here we report a *NPEPPS*-*TBC1D3* digenic genomic structure, which originated from joint SDs and became fixed in humans. The intact sequence of this digenic structure is absent in non-human primates. A human-specific hypomethylation signal within a CpG island pinpoints the expressed *TBC1D3* paralog that is transcribed predominantly as *NPEPPS*-*TBC1D3* readthrough transcripts. The readthrough transcripts underlie various post-transcriptional and translational regulation mechanisms, potentially fine-tuning the functions of TBC1D3. Moreover, readthrough transcription originating from joint SDs is potentially a broad mechanism for the emergence of new functions. Our work provides a necessary foundation for future functional studies and illustrates how joint SDs drive human-specific regulatory innovation.

## Discussion

Genomic reorganization resulting from large-scale structural variants is a prominent drive for functional innovation in evolution, often leading to the formation of new gene bodies or new regulatory mechanisms enabled by newly formed contact loops^5,28^. When a new gene is repositioned near another existing gene, the co-option of the neighbor’s regulatory elements may happen, reshaping the new gene’s expression pattern. This alteration may be a major step in the evolution of the new gene/transcript, potentially providing a wider expression range for further selection. Interestingly, this repositioning also creates a genomic structure that may serve as an “evolutionary doorway” to a readthrough transcript, which fine-tunes the new gene’s expression level through readthrough efficiency and splicing variability, as well as translation efficiency regulated by the proximal gene’s sequence as an untranslated region (UTR). Here, we describe the *NPEPPS*-*TBC1D3* digenic structure originating from a joint SD event as such an example.

There are growing interests in *TBC1D3* because of its uniqueness in primate evolution and its potential functions in the brain. Many exciting *TBC1D3* functions have been reported and proposed. For instance, heterologous expression of *TBC1D3* in cortical neural progenitor cells of developing mouse brains has been reported to result in a proliferation of outer radial glial cells and a cortical expansion and folding^23^, likely through suppressing the histone methyltransferase G9a^41^. Additionally, *TBC1D3* expression has been reported to promote dendritic arborization and delay synaptogenesis in pluripotent stem cell-derived human cortical neurons, potentially associated with neoteny^22^. Furthermore, some publications have claimed an association between *TBC1D3* and tissue repair^21^. However, previous studies substantially lack mechanistic insight into *TBC1D3* gene expression regulation. Hence, the discoveries in our study, especially the novel insights that *NPEPPS*-*TBC1D3* readthrough transcripts are the predominant source of *TBC1D3* expression in various cell types and the hypomethylated CpG island marks the active expression of the readthrough transcripts, bridge significant knowledge gaps in the multifaceted control of TBC1D3 expression.

In addition to the regulatory mechanisms explored above, *NPEPPS*-*TBC1D3* expression may also be regulated through other mechanisms. For instance, the poly(A) signal may play a role in transcription readthrough regulation^42^. The truncation of *NPEPPS*-39M eliminates the canonical poly(A) signal used by *NPEPPS*-48M, which potentially contributes to the 39M transcriptional readthrough (Supplementary Fig. 24). Additionally, a recent paper reported that capped *trans*-RNAs can initiate translation at its target site^43^. Although that study did not identify a *trans*-RNA for *NPEPPS*-*TBC1D3* in HEK293T cells, it would be worthwhile to investigate whether this mechanism operates in other, more relevant cell types.

Current studies of TBC1D3 protein functions largely rely on exogenous overexpression systems, and endogenous TBC1D3 peptides are only occasionally detected in proteomics studies, primarily in testis (Supplementary Fig. 25a). Notably, TBC1D3 expression was often undetectable, unidentified, or understudied in previous brain and brain organoid proteomics studies^44–49^. While one previous study did detect a TBC1D3 peptide DVVEVAGSWWAQER in brain and brain organoid samples^50^, its signal intensity was marginal (Supplementary Fig. 25b). Based on the regulatory mechanisms elucidated in our study, we conclude that investigating TBC1D3’s neurodevelopmental functions in a physiologically accurate manner necessitates supplementing current overexpression research with studies that account for intracellular protein levels and dynamics. This also calls for new paradigms in proteomics for studying proteins associated with complex genomic features.

The insights provided by our study are critical not only for understanding TBC1D3 functions in neural development but also for enhancing cancer research. *TBC1D3* was previously reported to be an oncogene which is “amplified” in prostate and breast cancers^51^. Following that, a growing number of studies have suggested that *TBC1D3* is overexpressed in various cancer types and could serve as a prognostic biomarker^52^. However, the extensive CNV observed in the general population (Fig. 1 and Supplementary Fig. 1,2) has rarely been taken into consideration. Meanwhile, the accurate pinpointing of the exact paralogs involved in the oncogenic process remains to be achieved through comprehensive genomic investigation and standardized annotations. Here, we showcased that the *NPEPPS*-*TBC1D3* readthrough transcripts may as well serve as the predominant source of TBC1D3 expression in certain cancer cells (Supplementary Fig. 26), which calls for extra attention in future cancer research.

Notably, in the *NPEPPS*-*TBC1D3* readthrough transcripts, while the *NPEPPS* sequence can be deemed as the 5’ UTR of *TBC1D3*, on the other hand, the *TBC1D3* sequence can be deemed as the 3’ UTR of *NPEPPS* as well. *NPEPPS* encodes the puromycin-sensitive aminopeptidase, implicated in peptide degradation, cell cycle control, and cell polarity^53–56^. Certain ORFs within the *NPEPPS* region in some of the readthrough transcripts overlap with the coding sequence of the annotated Peptidase M1 membrane alanine aminopeptidase domain (Pfam, PF01433; p.280-497, NP_006301) (Supplementary Fig. 27). Future study is required to determine whether these ORFs translate and how the *TBC1D3* sequence affects their regulation.

Finally, our analysis revealed the existence of 109 readthrough transcription gene pairs originating from joint SD pairs, suggesting a broad mechanism for the emergence of new functions. Future functional and biomedical investigations will be essential to further elucidate their biological significance.

## Methods

### CNVs and digenic structures of *NPEPPS* and *TBC1D3* in the human population

We identified populational CNVs of *NPEPPS* and *TBC1D3* by aligning complete gene segments to populational assemblies using minimap2 (v2.30) with the parameters ‘-cx asm20 --secondary=yes -p 0.001 -N 1000 --eqx -Y -K 8G -s 1000’. To resolve the genomic context of individual copies and assign them to distinct gene clusters, we employed the *NPEPPS*-48M (the ∼100-kbp full length copy) as the anchor. For each detected copy, we computed its relative genomic position with respect to this anchor and assigned it to a specific cluster based on the relative distance on T2T-CHM13. To detect digenic genomic structures in the human population, we aligned the 39M digenic genomic structure of T2T-CHM13 to the other assemblies and excluded the alignments that contained a large structural variant (>1 kbp) within the structure. To analyze the breakpoints of SDs, all identified gene copies are aligned to the *NPEPPS*-48M and *TBC1D3*-39M and samtools (v1.21) to visualize the alignment depths. All coordinates in the main text, if not otherwise stated, are according to T2T-CHM13.

### Isoform analysis and RNA-seq analysis

Public iso-seq data are aligned to human and NHP genomes respectively using pbmm2 with alignment prefix ‘--preset ISOSEQ --sort’. Jbrowse2 (v3.5.1) is used to visualize and analyze the splicing and readthrough of the *NPEPPS*-*TBC1D3* digenic structure. Cell samples for RNA-seq were snap frozen in RNA Extraction Solution (Servicebio, #G3013). RNA extraction, quality control, and Illumina NovaSeq sequencing were performed by Personalbio. We utilized the nf-core rnaseq pipeline^57^ to quantify the expression level of *TBC1D3* paralogs on both T2T-CHM13 and GRCh38 reference genome. To identify gene fusion/readthrough events on the whole genome using Iso-Seq data, we developed a tool called FusionSeekerPro (https://github.com/Zikun-Yang/FusionSeekerPro) to call fusion events and identify the breakpoints. Fusion events supported by fewer than one reads and gene pairs in opposite orientations are excluded. Gene expression levels were quantified from long read Iso-Seq data using oarfish (v0.9.0)^58^. We defined confident fusion events as those with ≥5 supporting reads and expression levels exceeding one. To assess the enrichment of JSDTR in the fusion events, we performed Fisher’s exact test followed by Benjamini-Hochberg correction for multiple testing. The background gene pair set was defined as all same-strand gene pairs within the genome whose intergenic distances fell within the 95th percentile of distances observed in our confident fusion events.

### Cell culture

Cells were cultured at 37 with 5% CO2. HEK293T cells (National Collection of Authenticated Cell Cultures, NCACC, #SCSP-502) were maintained in Dulbecco’s Modified Eagle’s Medium (DMEM; Servicebio, #G4515) supplemented with 10% fetal bovine serum (FBS; Servicebio, #G8003), and 1× Penicillin-Streptomycin (Servicebio, #G4003). SH-SY5Y cells (NCACC, #SCSP-5014) were maintained in DMEM/F-12 (Servicebio, #G4612) supplemented with 15% FBS, 1× Penicillin-Streptomycin. CN1 and C20 iPSC lines were established in Asian Pan-Genome Project Phase 1 (APGp1), and maintained in eTeSR (STEMCELL, #100-1215). For chimpanzee (*Pan troglodytes*) cell lines, blood samples were obtained from a 23-year-old female chimpanzee (Z2) and a 7-year-old female chimpanzee (D2) from the Shanghai Wild Animal Park colony in accordance with the corresponding IACUC protocol (No. SJTU-B2024001). Enriched chimpanzee peripheral blood mononuclear cells (PBMCs) were reprogrammed into induced pluripotent stem cells (iPSCs) under feeder-free conditions. NPCs were induced using STEMdiff SMADi Neural Induction Kit (STEMCELL, #08581) and maintained in STEMdiff Neural Progenitor Medium (STEMCELL, #05833). FBNs were induced using STEMdiff Forebrain Neuron Differentiation Kit (STEMCELL, #08600) and STEMdiff Forebrain Neuron Maturation Kit (STEMCELL, #08605).

### Generation of Chimpanzee iPSCs

PBMCs were cultured in StemSpan II medium (STEMCELL, #09605) for 9 days to expand blood progenitor cells. On the day of reprogramming (day 0), the cells were transfected with the Epi5 Episomal iPSC Reprogramming Kit (Thermo Fisher Scientific, #A15960) using a Neon NxT electroporation system (Thermo Fisher Scientific). The cells were then plated in 2 mL of StemSpan II medium in a 6-well plate coated with Matrigel (Corning, #354277). On day 2, 1 mL of StemSpan II medium was added to each well without removing the existing medium. On days 3 and 5, 1 mL of ReproTeSR medium (STEMCELL, #05926) was added to the existing medium. Starting on day 7, the culture medium was completely replaced with 2 mL of ReproTeSR medium daily until day 20, when the first iPSC colonies were observed. Each colony was manually picked using a 200 µL pipette and transferred to new Matrigel-coated plates in mTeSR Plus medium (STEMCELL, #100-0276,) supplemented with 2 mM thiazovivin (MCE, #HY-13257) on the first day after passage.

### 5’ Rapid Amplification of cDNA Ends (5’ RACE) and TOPO cloning

RNA was extracted from CN1 NPC and C20 NPC using the FastPure Complex Tissue/Cell Total RNA Isolation Kit (Vazyme, #RC113). 5’ RACE was performed with the HiScript-TS 5’/3’ RACE Kit (Vazyme, #RA101). Two *TBC1D3* Gene Specific Primers (GSPs) were utilized in this study: 5’-TTCGAAAAGGCTTAGGCCCCTTGTCC-3’ (*TBC1D3*-Rev0) and 5’-CAGCTCCGTCTCATGTACAATCCCCAAA-3’ (*TBC1D3*-Rev1). 5’ RACE products were purified with the FastPure Gel DNA Extraction Mini Kit (Vazyme, #DC301) and utilized for TOPO cloning using the Ultra-Universal TOPO Cloning Kit (Vazyme, #C603). Subsequently, the TOPO cloning reaction was transformed into the Fast-T1 competent cells (Vazyme, #C505). Ten colonies from each 5’ RACE reaction of each cell line were picked for Sanger sequencing using the corresponding 5’ RACE GSP as the sequencing primer. Additionally, the 5’ RACE products were also analyzed using Nanopore sequencing (GenScript).

### Single-cell analysis on human brain organoids

Single-cell transcriptomic profiling was carried out on two previously established brain organoid lines (CN1 and C50) collected at differentiation days 42 and 98 (Han *et al.*, manuscript in preparation). Libraries were constructed with the 10× Genomics Chromium system. Sequencing reads were mapped to the T2T-CHM13v2.0 (GCF_009914755.1-RS_2023_10)^12^ and cell-by-gene count matrices were produced using Cell Ranger (v7.2.0)^59^. Standard quality-control filters were applied to exclude low-quality cells and genes following recommended practices^60^. Putative doublets were identified and removed using DoubletDetection (v4.2)^61^. Preprocessed datasets were integrated, and batch effects were corrected with harmonypy (v0.0.10)^62,63^. Downstream analysis was performed using Scanpy (v1.11.0)^64^ and cell type annotations were assigned manually using canonical marker genes.

### Evolutionary history reconstruction in African great apes

Human chromosome 17 and its nonhuman primate counterparts are aligned using minimap2, with secondary alignment allowed, and visualized by SVbyEye (v0.99.0). Estimated copy numbers of *TBC1D3* and *NPEPPS* in primates are calculated by fastCN. To characterize the *NPEPPS-CCL4* in bonobos and chimpanzees, Iso-Seq data were first examined, which indicated that readthrough transcripts spanning *NPEPPS* and *CCL4* represent the predominant isoforms in both species. The readthrough ratio is estimated as the proportion of readthrough-supporting split-read depth relative to the total read depth at a representative genomic position. For bonobo, the position with the maximal accumulation of readthrough-supporting split-reads was identified and used to estimate the readthrough ratio. However, when readthrough ratios were estimated similarly for chimpanzees, the ratios were substantially lower than expected based on Iso-Seq results. To investigate this discrepancy, bonobo readthrough-supporting split-reads and the genomic sequence of the bonobo *CCL4* locus (*LOC100991964*) were independently aligned to the chimpanzee genome. These cross-species mappings revealed that the *CCL4* sequence of the bonobo *NPEPPS*-*CCL4* splice junction corresponds to the terminal annotated exon of *NPEPPS* in chimpanzees, indicating differences in gene annotations between the two species. Based on the Iso-Seq evidence and the cross-species genomic alignment, we inferred that the direct estimation of the readthrough ratio from chimpanzee bulk RNA-seq data underestimates the true abundance of *NPEPPS-CCL4* readthrough transcripts among all *CCL4* transcripts. Therefore, to obtain a more accurate estimate of the readthrough ratio in chimpanzees, the orthologous position to the bonobo representative position was determined in the chimpanzee genome and used to estimate the ratio.

### Ribo-Seq analysis

*TBC1D3*-only sequences and *NPEPPS*-*TBC1D3* sequences were cloned from SH-SY5Y cDNA, which was synthesized as described above. 5’-ATGGACGTGGTAGAGGTCGC-3’ and 5’-CTAGAAGCCTGGAGGGAACTGAG-3’ were used as the matching sequences of the primers to amplify *TBC1D3*-only sequences. 5’-CTAGAAGCCTGGAGGGAACTGAG-3’ (the reverse primer), 5’-CGCCTCCTTCCCAACCCC-3’(the forward primer for NT1-1 and NT1-2), and 5’-ATGAATTGTGCTGATATTGATATTATTACAGC-3’ (the forward primer for NT2-1, NT2-3, NT2-4, and NT2-5). PCR reactions were performed with Q5 High-Fidelity DNA Polymerase (NEB, #M0491) and assembly reactions were performed with NEBuilder HiFi DNA Assembly Master Mix (NEB, #E2621), following the manuals. 2.1∼2.6 ug of each plasmid was transfected to HEK293T plated in a 6-well plate well using Lipomaster 3000 (Vazyme, #TL301), ensuring the same amount of molecules was used. Cells were treated with cycloheximide (final conc. 0.1 mg / mL; Selleck, #S7418) and sent for Ribo-seq (Novogene) on Day 4 post-transfection. bowtie2 (v2.5.4) was utilized to map the sequencing reads to the rRNA sequences from T2T-CHM13v2.0 in order to achieve the target sequences with the parameter ‘--un-gz’. Then the cleaned reads were aligned to the manual annotated sequences and T2T-CHM13v2.0 using bowtie2.

### Immunoblotting

Cells were washed once with PBS (Servicebio, #G4202) and lysed in RIPA buffer (Sigma-Aldrich, #R0278) supplemented with cOmplete Protease Inhibitor Cocktail (Roche, #04693116001). The lysates were incubated on ice on a shaker for 15 min, followed by centrifugation to collect the supernatant (16,000 × g for 30 min at 4 °C). Protein concentration was determined using the BCA Protein Quantitative Detection Kit (Servicebio, #G2026). A total of 15 μg of protein per sample was loaded onto an 10% SDS–PAGE gel (Genefist, #GF1820-10). After electrophoresis, proteins were transferred onto a PVDF membrane (Servicebio, #G6044-0.45) and blocked for 25 min with Protein Free Rapid Blocking Buffer (Servicebio, #G2052). The membrane was washed with TBST and incubated overnight at 4 °C on a shaker with the following primary antibodies: GFP (abcam, #ab290, 1:5000), ACTB (HUABIO, #EM21002, 1:20000), and TBC1D3 (Santa Cruz Biotechnology, #sc-376073, 1:100). After washing with TBST, the membrane was incubated with HRP-labeled goat anti-mouse IgG (Servicebio, #GB23301, 1:5000) for 1 h at room temperature on a shaker, and then washed again. The protein bands were imaged using Clinx ChemiScope 6000.

## Data Availability

### Genome assemblies

The T2T primate genomes used in this study are available from GenBank via accessions: GCA_009914755.4, GCA_028858775.2, GCA_028885625.2, GCA_028885655.2, GCA_029281585.2, GCA_029289425.2 and GCA_037993035.1. The T2T primate genome assemblies are also available on GitHub (https://github.com/marbl/Primates and https://github.com/zhang-shilong/T2T-MFA8). We used the human assemblies from Human Pangenome Reference Consortium (HPRC), Human Genome Structural Variation Consortium (HGSVC) and Asian Pan-Genome Project (APG).

### Transcriptome data

Iso-seq generated from the human CHM13hTERT cell line can be accessed by SRR12519035 and SRR12519036. Nonhuman primate Iso-seq data are deposited under BioProject identifiers PRJNA902025, PRJNA1016395 and PRJNA1041301. Raw bulk RNA-seq data for chimpanzee, gorilla, and macaque tissues were obtained from published studies^65–68^ (PRJNA1004471, PRJNA143627, PRJNA304995, and PRJNA236446).

### Other data

Other data generated in this study, including the RNA-seq results of the CN1 cell lines and the chimpanzee iPSCs, the 5’ RACE sequencing data, the Iso-Seq data of CN1 and C20 NPCs, and the Ribo-Seq data, are available on request from the corresponding authors following the regulatory guidelines. Single-cell RNA-seq data for organoid lines CN1 and C50 at differentiation days 42 and 98 are presented in Han *et al.* (manuscript in preparation) and are available from the authors on request.

## Code Availability

The custom scripts used in this study are available on GitHub (https://github.com/YafeiMaoLab/TBC1D3_analysis).

## Acknowledgements

We would like to acknowledge the Human Pangenome Reference Consortium (BioProject ID: PRJNA730823), the National Human Genome Research Institute (NHGRI), Human Genome Structural Variation Consortium (HGSVC), Asian Pan-genome project (APG), and Primate T2T Consortium for providing long-read human and great ape genome assemblies. We thank Dr. Lin Ge (Beijing Children’s Hospital) for the suggestions regarding SMA. This work is supported by the Shanghai Rising-Star Program for studying “evolutionary medicine and functional studies of the human-specific *NPEPPS* duplications” (24YF2721800 to K.M.). This work is in part supported by the Shanghai Magnolia Talent Plan Pujiang Project (24PJA049) and the Shanghai Post-doctoral Excellence Program (2024338) to K.M., and by Shanghai Jiao Tong University Medical-Engineering Interdisciplinary Research Fund (grant no. YG2025QNA47) to X.Y. This work is in part supported by the National Natural Science Foundation of China (32130035) to Z.-G. L. This work is in part supported by the Scientific Research Innovation Capability Support Project for Young Faculty (SRICSPYF-ZY2025101), the National Key Research and Development Program of China (2025YFC3410300), the National Natural Science Foundation of China (32370658), the Natural Science Foundation of Chongqing, China (CSTB2024NSCQ-JQX0004), the New Cornerstone Science Foundation through the XPLORER PRIZE, Shanghai Jiao Tong University (SJTU) 2030 Initiative (WH510363003/016), Yongxin Youth Award Fund and Zhongying Young Scholars Program to Y.M.

## Author Information

### Contributions

K.M., Z.Y., and Y.M. conceptualized the study. K.M., Z.Y., and Z.L. contributed equally in terms of invested time and efforts, and therefore each has the right to list their name first in their respective curriculum vitae; we petition the grant funding bodies to treat them as true equals. K.M., Z.Y., Z.L., J.G., D.L., Z.W., H.M., S.Z., L.F., H.L., and X.J. performed the 5’ RACE and the transcription/translation analyses. Z.Y., K.M., Z.L., J.G., S.Z., L.F., G.L., J.C., J.Z., X.Y., G.Z., and Y.M. performed the population, evolution, and expression analyses. Z.L., K.M., Z.W., Q.X, C.Y., and Z.-G. L. conducted the functional studies. X.Y. generated the iPSCs, with P.L.’s assistance. Z.Y. developed the FusionSeekerPro tool. G.Z. initiated the APG project. K.M., Z.Y., and Z.L. wrote the original draft of the manuscript. K.M. and Y.M. edited the manuscript. All authors reviewed the manuscript.

### Corresponding authors

Correspondence to Kaiyue Ma and Yafei Mao

