## Supplementary Figures for "Joint Segmental Duplication Co-option Drives Human-specific Transcriptional Readthrough and Expression Fine-tuning of *NPEPPS*-*TBC1D3*"

Fig. S1: The haplotypes of 37M, 39M, 48M, 61M and 63M in the African population (n=68 haplotypes).

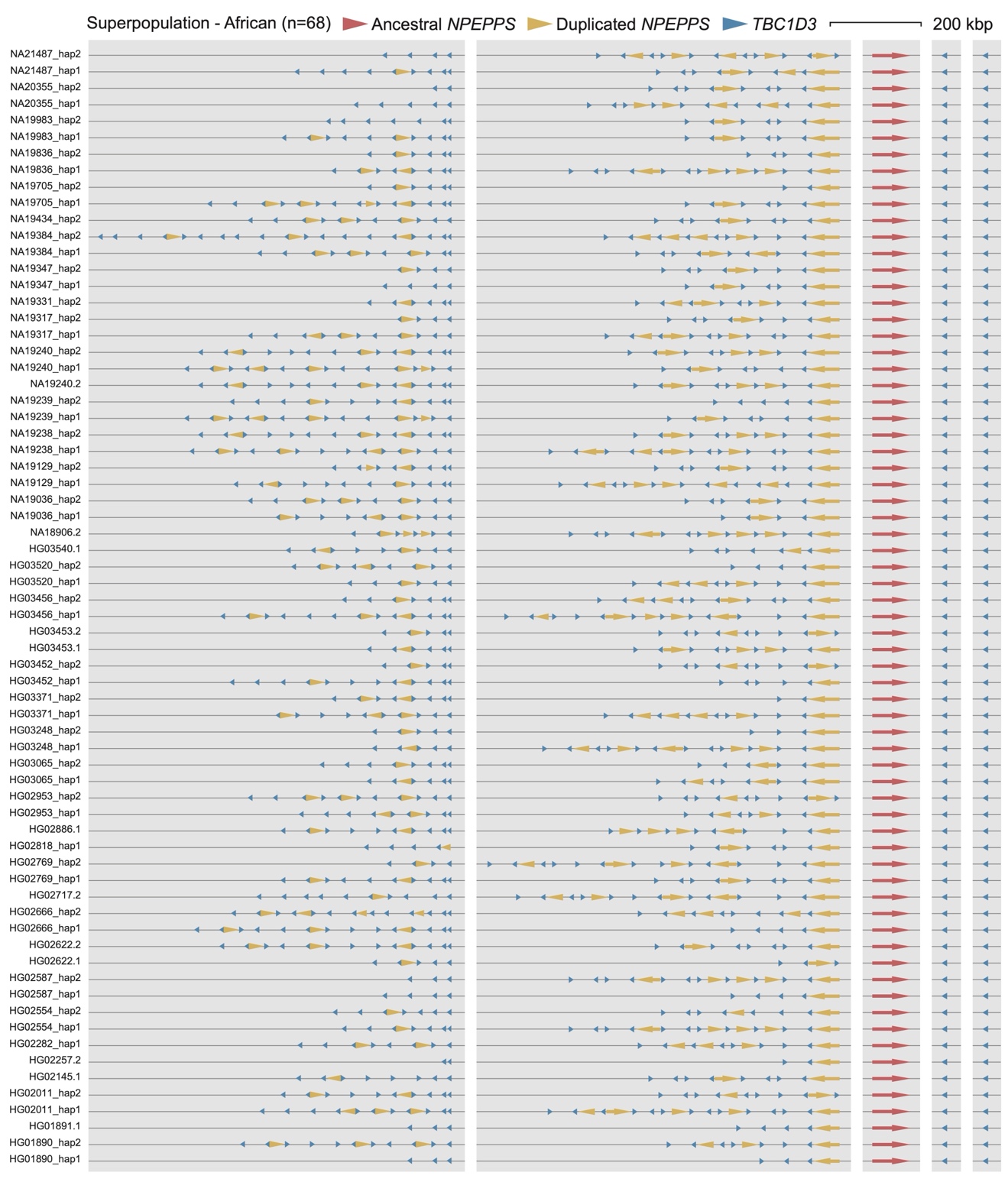

The haplotypes of 37M, 39M, 48M, 61M and 63M in the African population (n=68 haplotypes). The red, yellow, blue triangles represent the ancestral *NPEPPS*, duplicated *NPEPPS* and *TBC1D3* copies, respectively.

Fig. S2: The haplotypes of 37M, 39M, 48M, 61M and 63M in non-African populations (n=393 haplotypes).

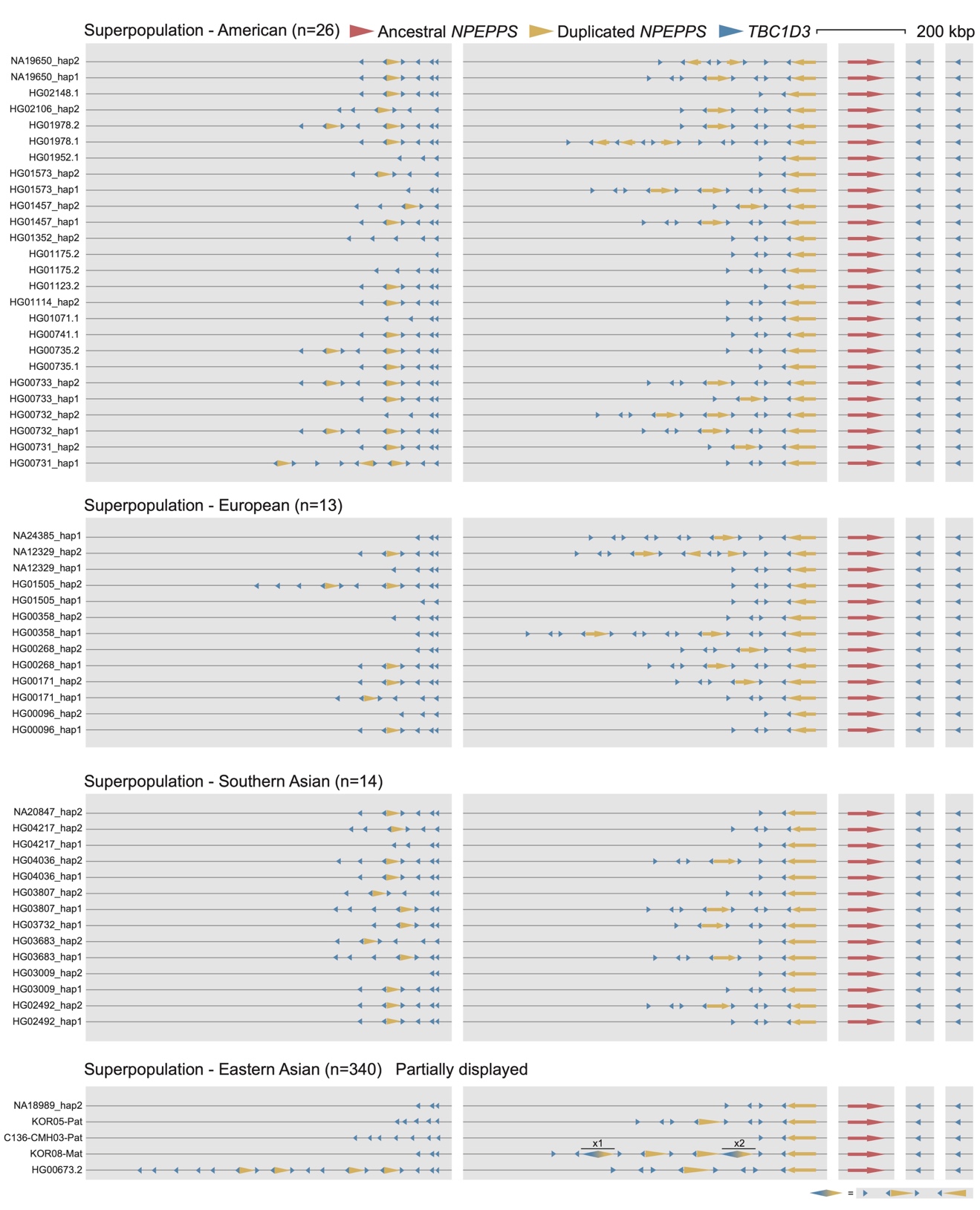

The haplotype of 37M, 39M, 48M, 61M and 63M in non-African populations (n=393 haplotypes). The red, yellow, blue triangles represent the ancestral *NPEPPS*, duplicated *NPEPPS* and *TBC1D3* copies, respectively. Haplotypes of Eastern Asian are partially displayed.

Fig. S3: The length and copy number variation for *NPEPPS* and *TBC1D3* segmental duplications.

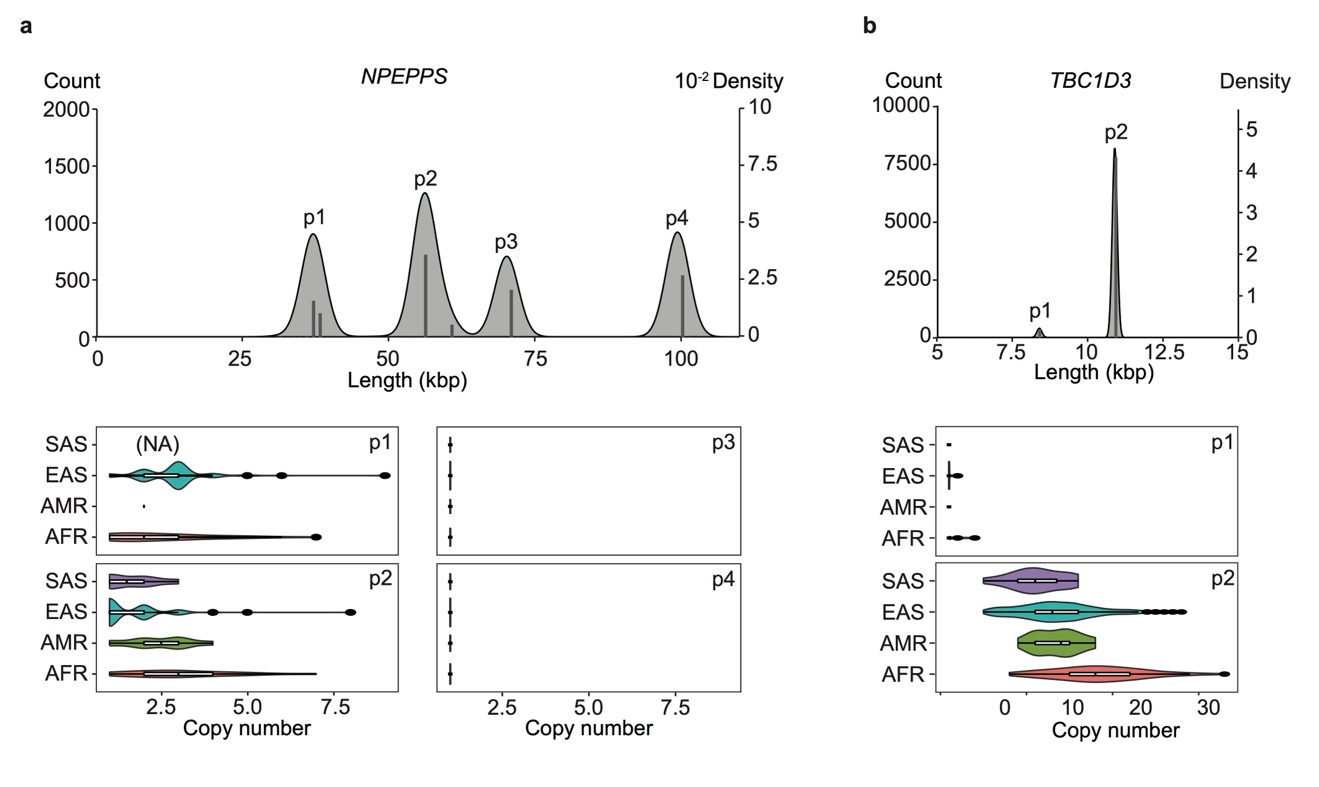

**a**, The length distribution of *NPEPPS* segmental duplications indicates incomplete duplicates of *NPEPPS* enriched at three peaks (p1, p2, p3). The bottom panel reveals the constrained copy number of p3 (corresponding to the NPEPPS incomplete duplicate that retains the hypomethylated CpG island) and p4 (the ancestral locus). **b**, The length distribution of *TBC1D3* segmental duplications indicates most of the *TBC1D3* copies are complete duplicates and contribute to the major part of the copy number variations.

Fig. S4: Previous attribution of *TBC1D3* expression to a specific *TBC1D3* paralog is largely influenced by the genome assembly and the gene annotation.

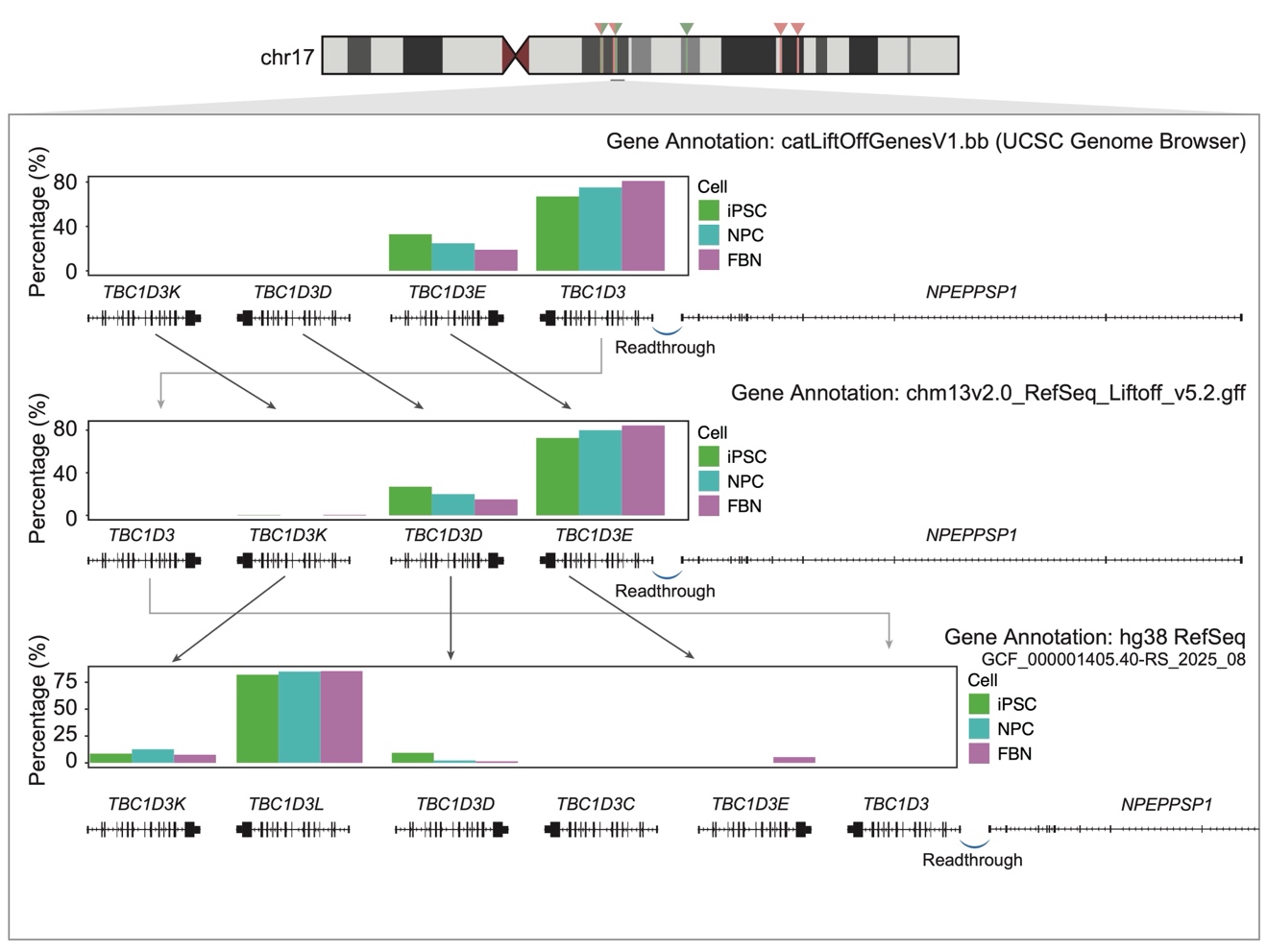

STAR and RSEM were used to quantify the *TBC1D3* expression attributed to different paralogs. The reference assembly and gene annotation versions substantially affect the results, potentially impacting the reliability of subsequent functional studies. To illustrate the inconsistencies in gene annotation across the same or different genomes, we retrieved the RefSeq annotations from multiple sources: UCSC genome browser[^1^](https://sciwheel.com/work/citation?ids=69929&pre=&suf=&sa=0), latest v5.2 RefSeq annotation (https://github.com/marbl/CHM13) and RefSeq annotation of GRCh38. STAR (v2.7.11b) and salmon (v1.10.3) are applied to align and quantify. Gene expression abundance was measured in terms of Transcripts Per Million (TPM). To identify the predominantly expressed paralogs in humans, we aggregated all the paralogs except *TBC1D3*-61M (*TBC1D3P1*) and *TBC1D3*-63M (*TBC1D3P2*) and calculated the proportion.

Fig. S5: Location, expression and disease association analysis of *TBC1D3* pseudogenes.

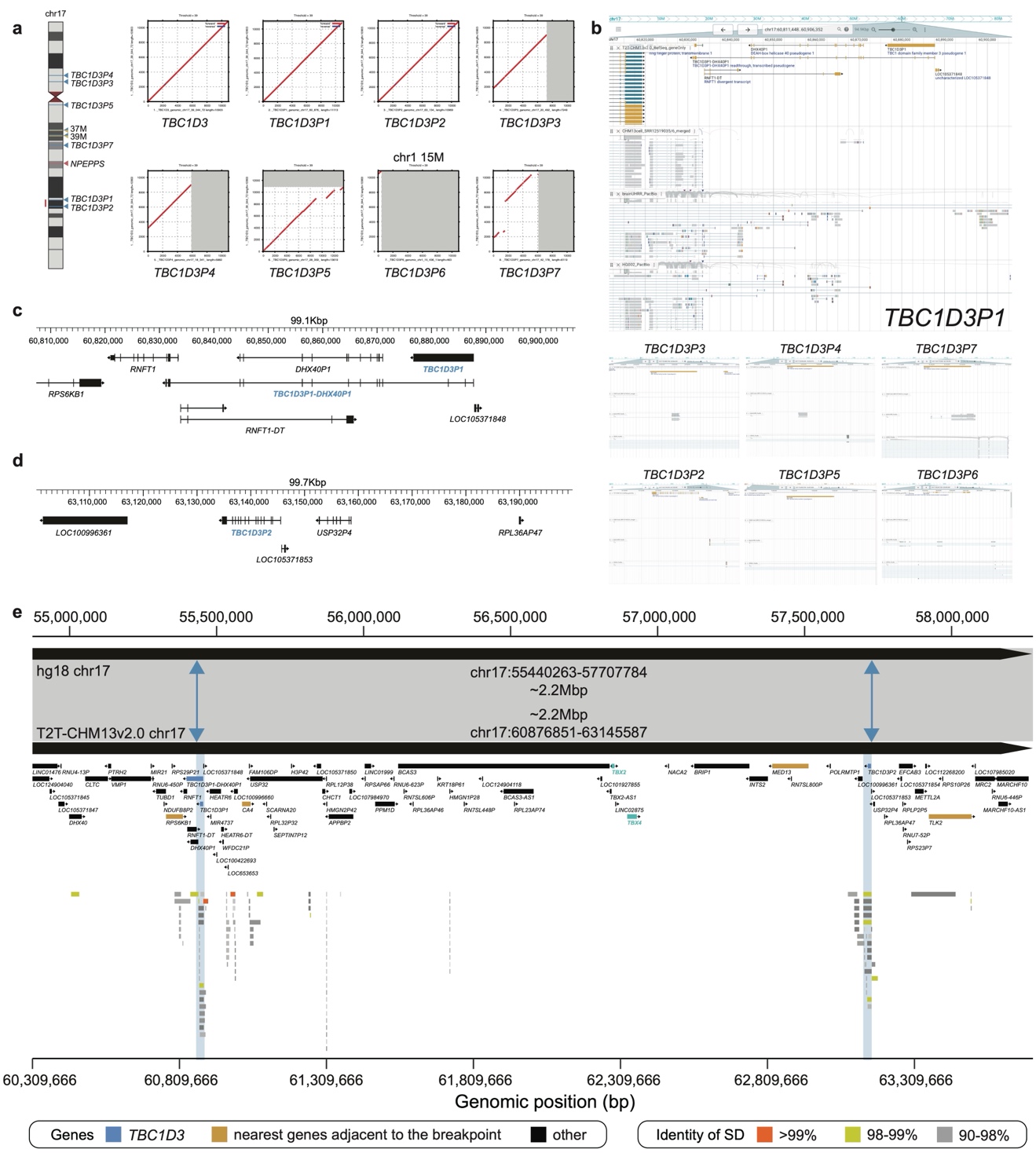

**a**, G-banding diagram of human chromosome 17 with highlighted locations of *TBC1D3* and *NPEPPS* copies. Pairwise alignments between *TBC1D3*-39M and *TBC1D3* pseudogenes indicate that *TBC1D3P1* and *TBC1D3P2* are nearly complete pseudogenes. **b**, Iso-Seq alignment tracks of the *TBC1D3* pseudogenes in CHM13hTERT cells and human brain tissues exhibit no read supporting their expression, excluding the occasional, potentially misaligned reads. **c**, Gene annotation of the *TBC1D3P1* locus. **d**, Gene annotation of the *TBC1D3P2* locus. **e**, The synteny comparison between hg18 and T2T-CHM13v2.0 shows the recurrent ~2.2-Mbp deletion in 17q23.1-q23.2 is mediated by the *TBC1D3P1* and *TBC1D3P2* segmental duplications.

Fig. S6: *NPEPPS*-*TBC1D3* readthrough transcripts naturally occur across diverse cell types in healthy individuals.

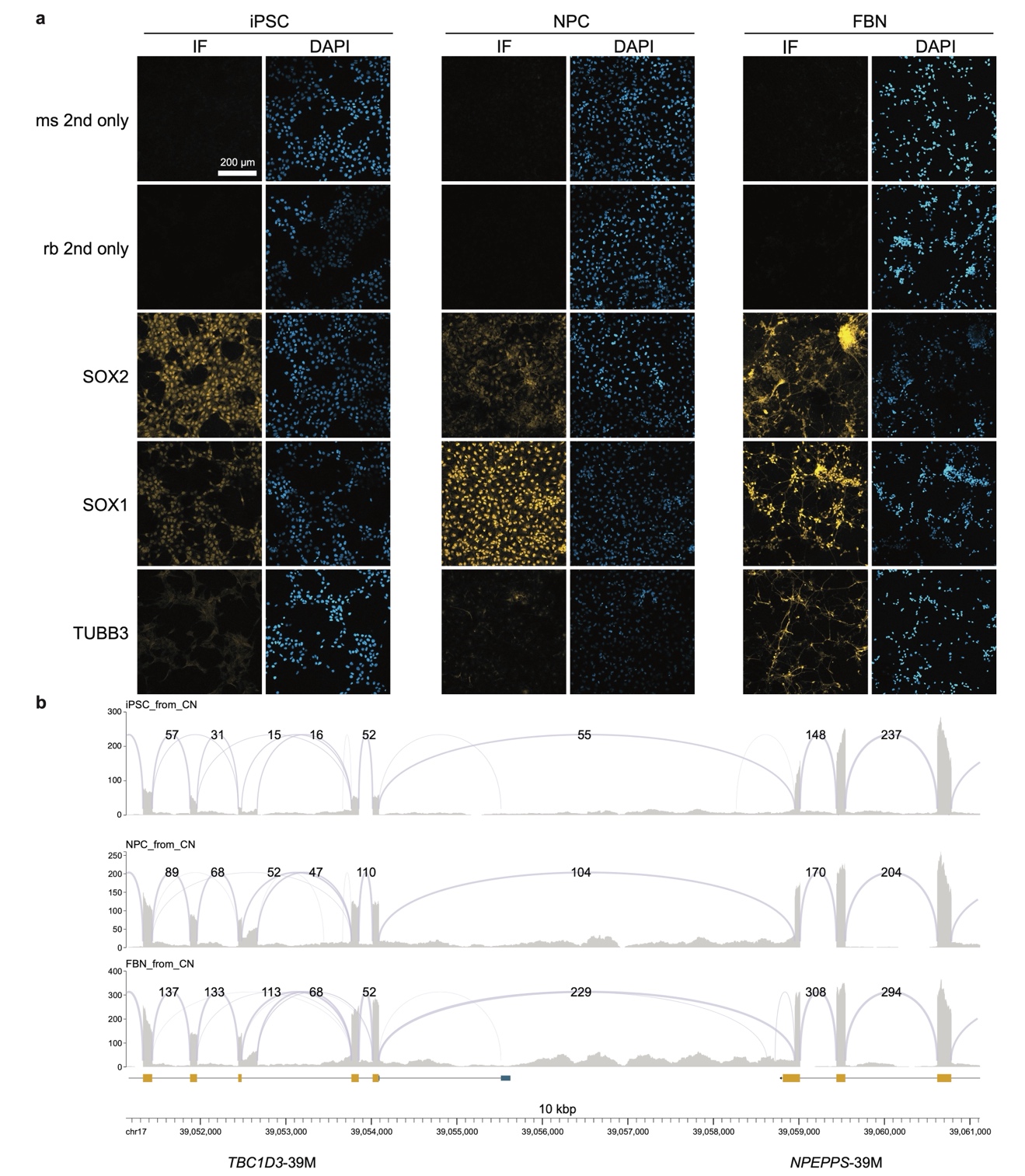

**a**, Cells plated on coverslips were fixed at 4 °C for 30 min with Paraformaldehyde Fixative (Neutral) (Servicebio, #G1101), permeabilized with 0.3% Triton X-100 (Solarbio, #T8200) in PBS at 4 °C for 30 min, and blocked with 5% Fetal Bovine Serum (FBS) in PBS at room temperature for 1 h. The cells were then incubated overnight with the following primary antibodies in PBS containing 1% FBS at 4 °C with gentle shaking: SOX1 (Invitrogen, #MA5-32447, 1:200), SOX2 (CST, #4900, 1:200), and TUBB3 (CST, #5568, 1:200). After washing with PBS, the cells were blocked using 5% FBS at room temperature for 1 h, and then incubated with corresponding secondary antibodies for 2 h at room temperature (Servicebio, #GB27301 and #GB27303, 1:500). Coverslips were then mounted on glass slides using Anti-fade Mounting Medium (with DAPI) (Servicebio, #G1407) and stored at 4 °C for the mounting medium to dry. Imaging was performed using Zeiss LSM 900. **b**, Splicing profiles of iPSC, NPC, FBN transcriptomes demonstrate universal existence of *NPEPPS*-*TBC1D3* readthrough transcript in diverse cell types.

Fig. S7: 5’ RACE-TOPO cloning reveals *NPEPPS*-*TBC1D3* readthrough transcripts as the predominant source of *TBC1D3* expression in NPCs.

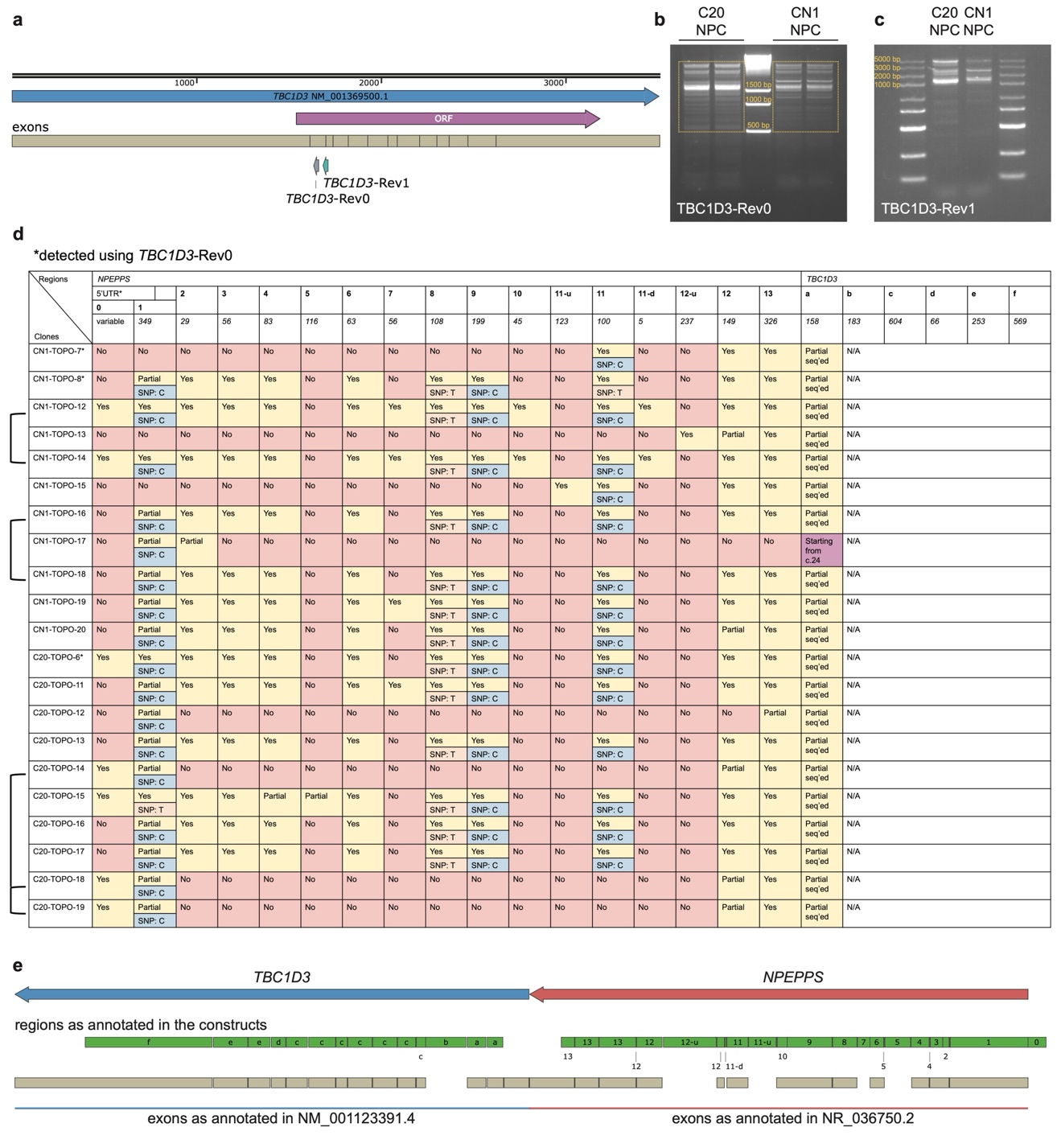

**a**, Two gene-specific primers were designed for 5’ RACE. *TBC1D3*-Rev0 was used in the pilot experiment. However, potentially because its sequence resides entirely within a single exon, the 5’ RACE generates many non-specific products and products amplified from gDNA. Therefore, *TBC1D3*-Rev1, which spans an exon junction, was designed for the optimized 5’ RACE experiments. **b**, The PCR products of the pilot 5’ RACE using *TBC1D3*-Rev0 were purified through gel purification (rectangle: the gel cut for purification). To achieve a higher recovery and a more accurate product representation, we used PCR purification for the *TBC1D3*-Rev1 5’ RACE. **c**, PCR products of the *TBC1D3*-Rev1 5’ RACE. **d**, TOPO cloning followed by Sanger sequencing detected different readthrough transcripts (rows connected on the left are those colonies with the same readthrough transcripts). The changing blocks in the sequences were labeled with numbers for *NPEPPS* and letters for *TBC1D3*. 5’ UTR is defined according to: NM_006310.4. Partial: only partial sequence exists. Partial Seq’ed: only a part of the region was covered by 5’ RACE, leaving the intactness of the region unknown. N/A: sequence existence is unknown due to primer design. The alternate SNPs are all only observed in one case. It can not be ruled out that they may be PCR artifacts. Note: 10 colonies were sequenced per GSP per cell line (40 in total). Only those definitely resolved are listed here (Rev0: 3; Rev1: 18). Colonies that originated from gDNA contamination (Rev0: 3), or non-specific amplification (Rev0: 7; Rev1: 1), or those generated noisy Sanger sequencing results (Rev0: 5; Rev1: 1), or those that were not definitely resolved (Rev0: 2), were excluded from this list. C20-TOPO-2 contains a *H2AZ1* sequence and C20-TOPO-7 was not fully resolved after all DNA samples were used for multiple Sanger sequencing attempts. Therefore, these two are not listed here. **e**, The labelled blocks largely overlap with annotated exons, indicating that the different transcripts arise from different splicing events.

Fig. S8: *NPEPPS*-*TBC1D3* readthrough transcript sequences reconstructed from C20 NPC 5’ RACE-TOPO cloning (Part 1).

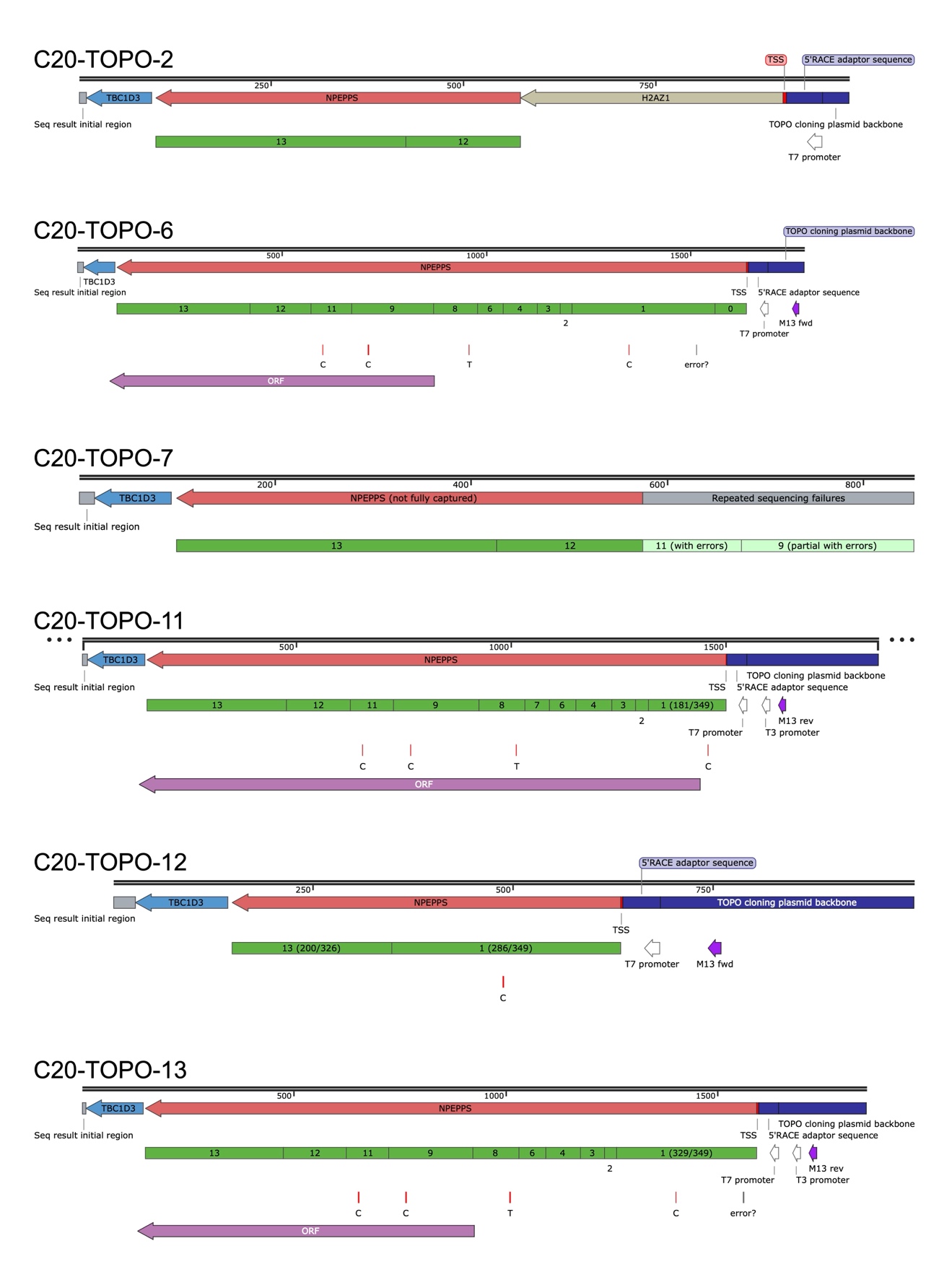

C20-TOPO-2 contains a *H2AZ1* sequence. It is unclear whether this transcript indeed exists, either in normal cells or in a cell carrying a structural variation. Alternatively, this may be an artifact introduced during the 5’ RACE-TOPO cloning process. C20-TOPO-7 was not fully resolved after all DNA samples were used for multiple Sanger sequencing attempts. Only ORFs with a size larger than 200 AA are shown (same in Supplementary Fig. 9-11).

Fig. S9: *NPEPPS*-*TBC1D3* readthrough transcript sequences reconstructed from C20 NPC 5’ RACE-TOPO cloning (Part 2).

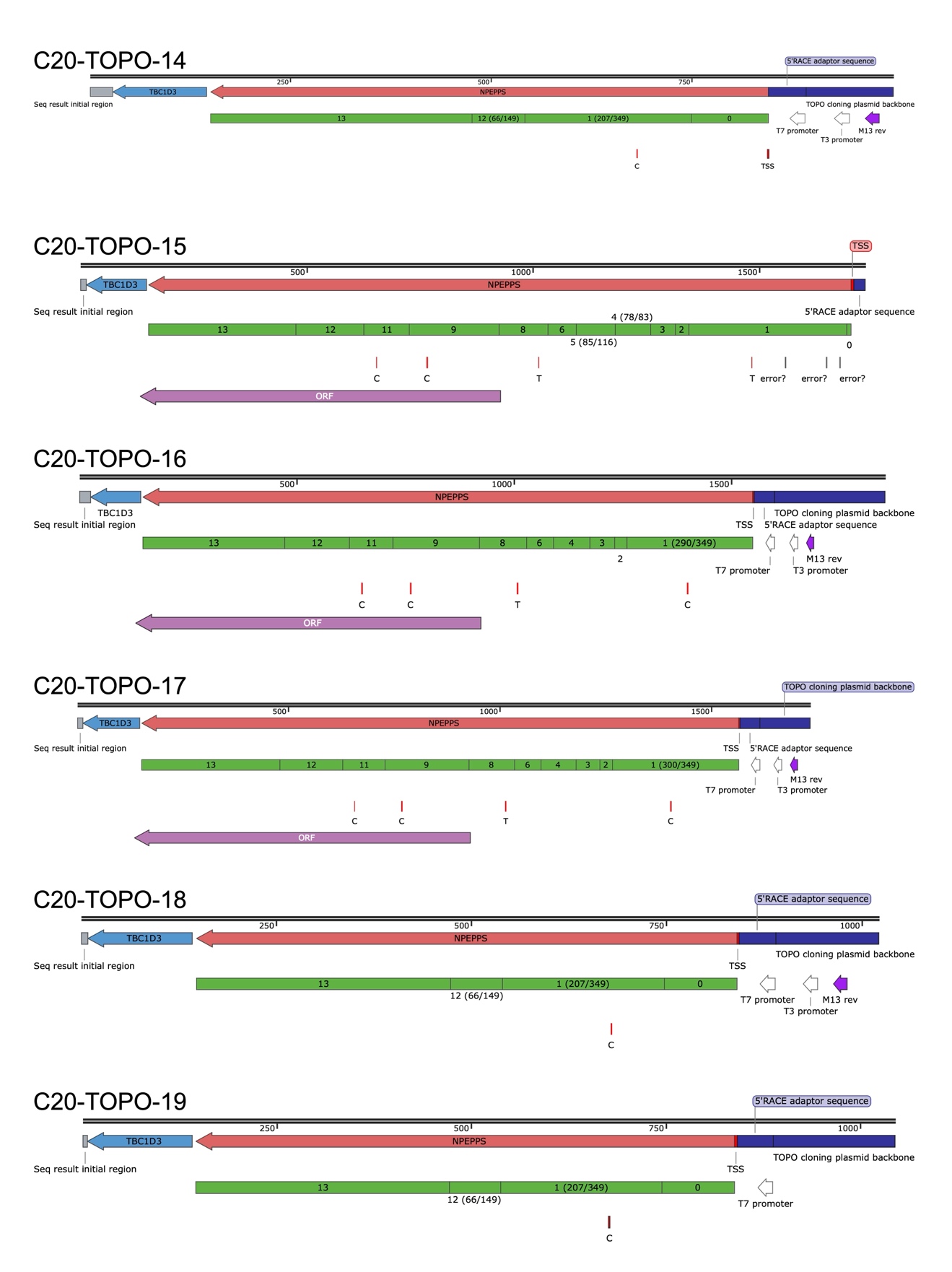

Fig. S10: *NPEPPS*-*TBC1D3* readthrough transcript sequences reconstructed from CN1 NPC 5’ RACE-TOPO cloning (Part 1).

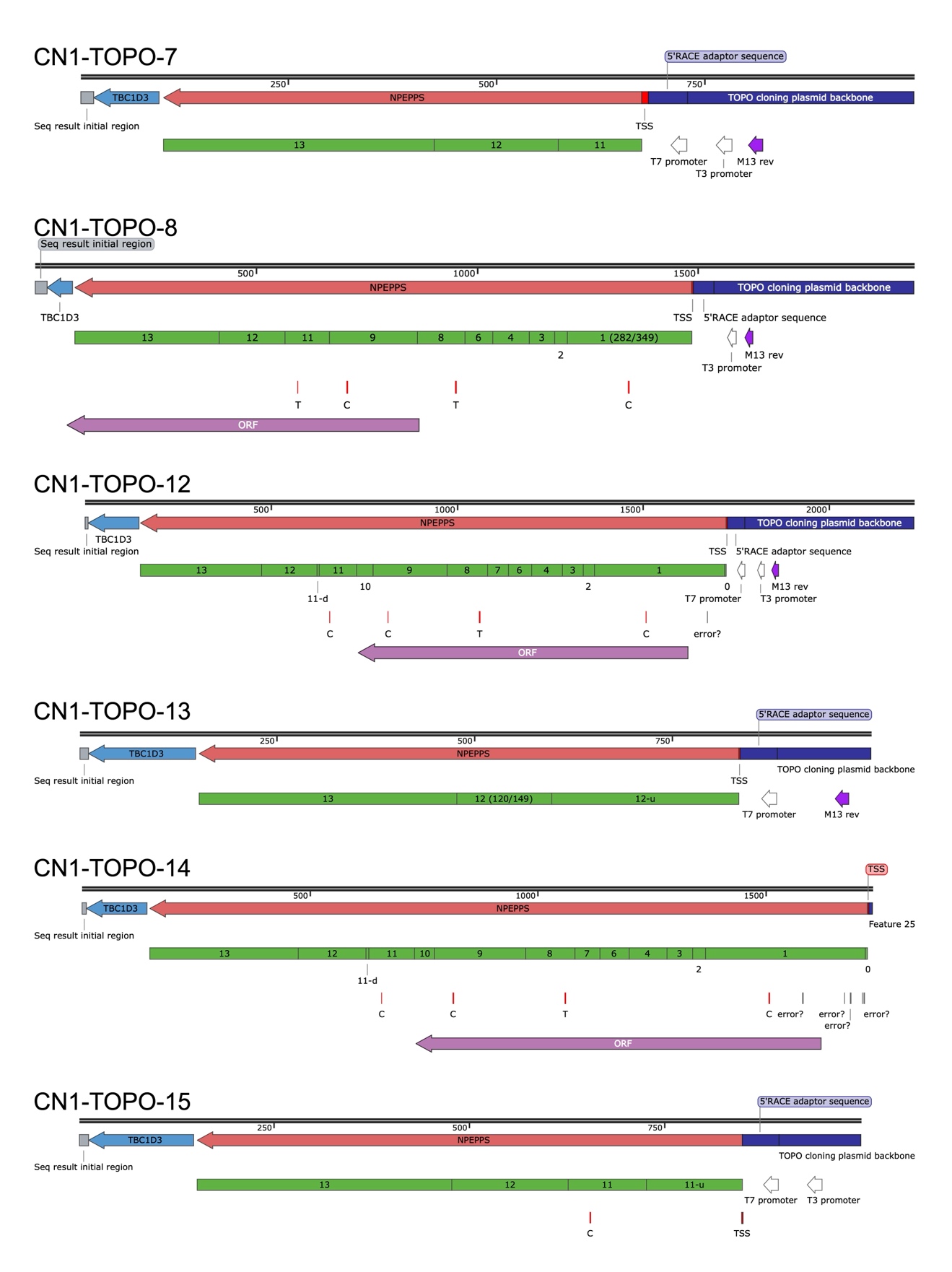

Fig. S11: *NPEPPS*-*TBC1D3* readthrough transcript sequences reconstructed from CN1 NPC 5’ RACE-TOPO cloning (Part 2).

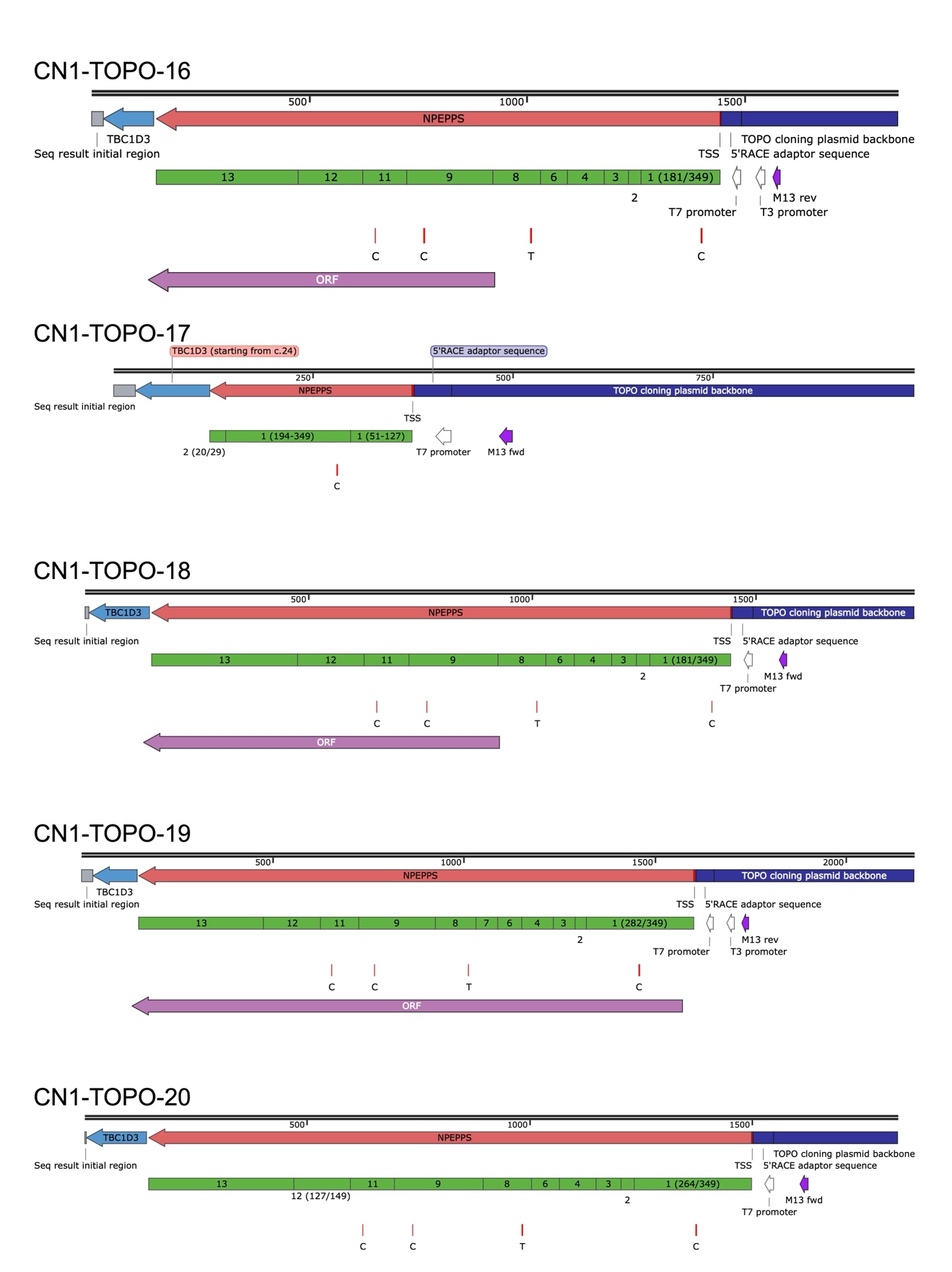

Fig. S12: 5’ RACE-nanopore processing and readthrough verification across three cell types.

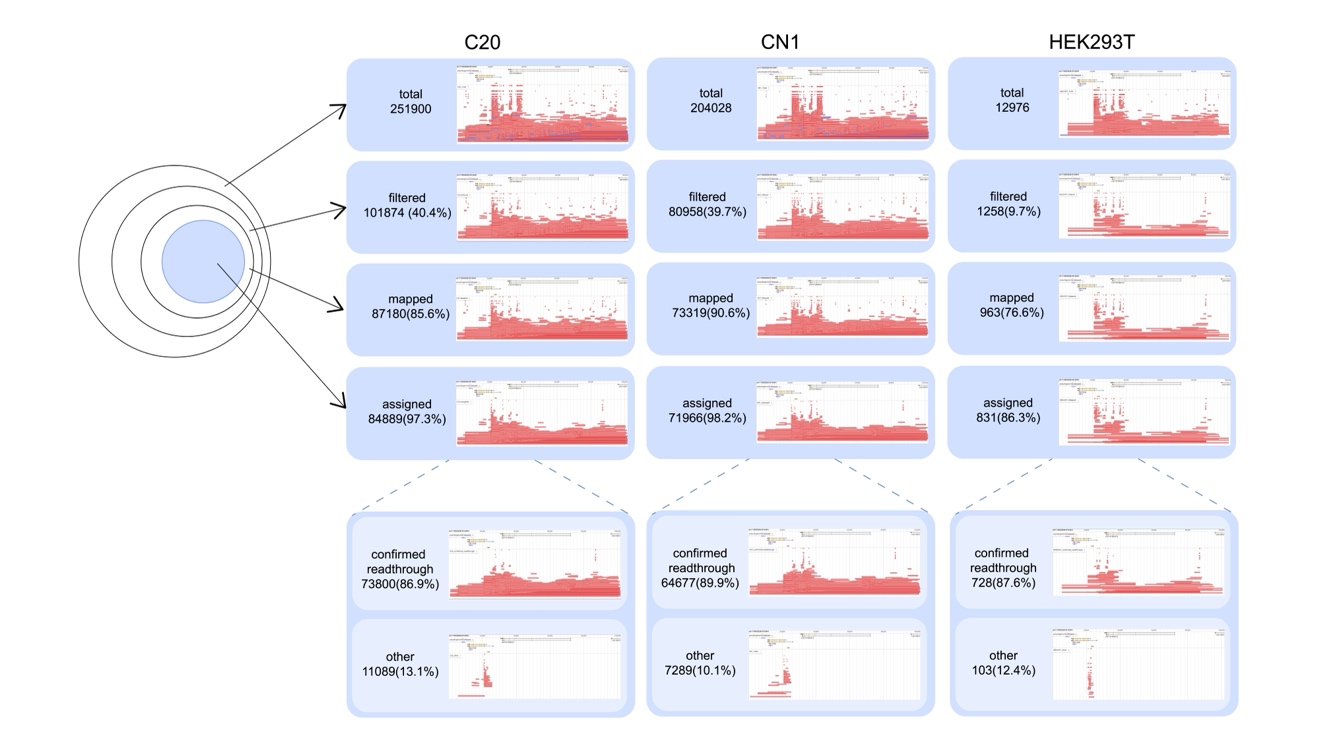

Stepwise 5’ RACE read processing in C20 NPC, CN1 NPC, and HEK293T cells. Read counts, retention rates, and JBrowse2 read-cloud display are shown across the sequential filtering steps listed as total, filtered, mapped to genome, assigned, and confirmed-readthrough/other. The split-read connections at the *NPEPPS*-*TBC1D3* junction reveal the predominance of readthrough transcripts.

Fig. S13: *NPEPPS*-*TBC1D3* readthrough quantification across iPSC-derived lineages and healthy testicular tissues.

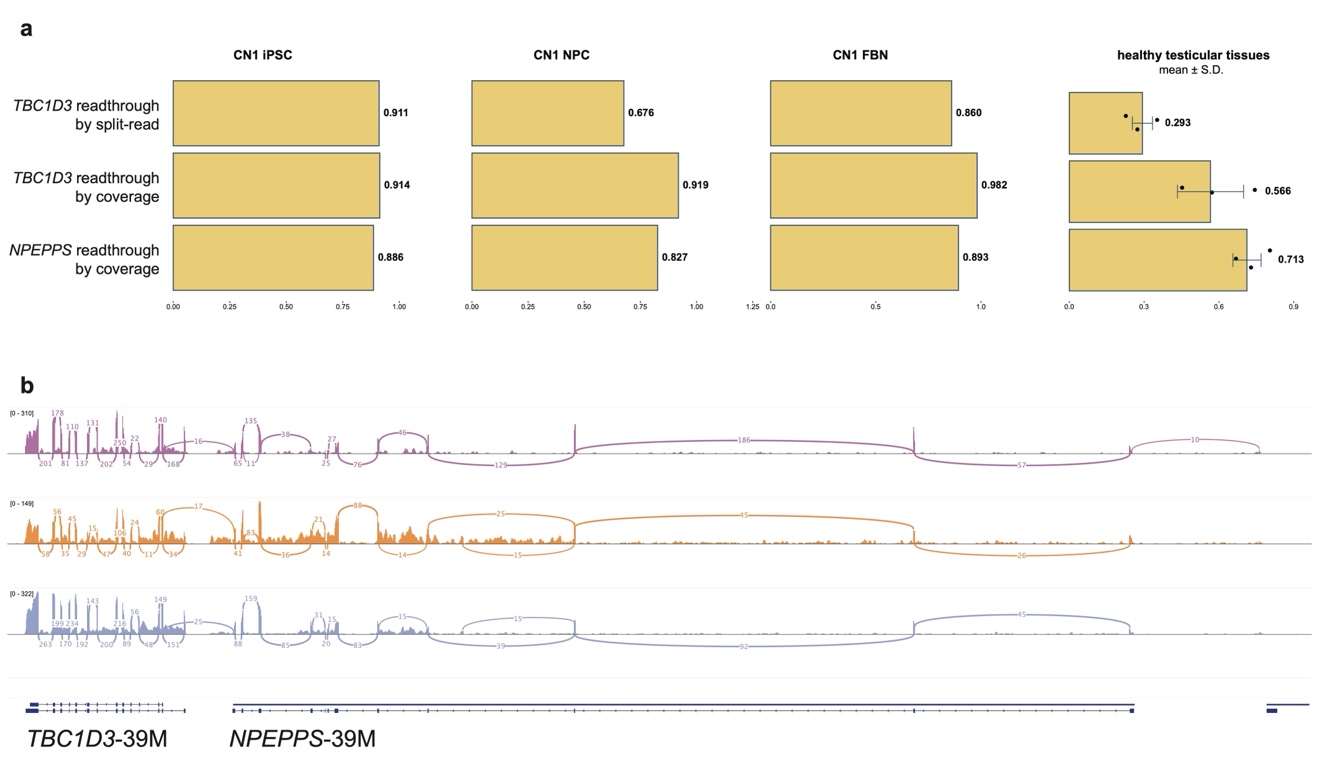

**a**, Quantification of *NPEPPS*-*TBC1D3* readthrough across CN1 cell lines (iPSC, NPC, FBN) and healthy testicular tissues. The bar plots display three readthrough metrics for each sample: (1) *TBC1D3* readthrough ratio supported by split-read evidence; (2) *TBC1D3* readthrough inferred from read-depth coverage; (3) *NPEPPS* readthrough inferred from read-depth coverage. **b**, The sashimi plots illustrate the read coverage and splicing junctions at 39M locus in healthy testicular tissues across three RNA-seq replicates. Note: Due to the limitations of short-read analyses, all the ratios should only be considered as rough estimations. (1) Split-read-based ratio for *TBC1D3*: the genomic position covered by the maximal number of readthrough-supporting split-reads was identified, and the readthrough ratio was defined as the proportion of the readthrough-supporting depth over the total depth at this position. (2) Coverage-based ratio for *TBC1D3*: as the last exon of *NPEPPS* partially skips to exon 2 of *TBC1D3*, exon 1 of *TBC1D3* are assumed to reflect non-readthrough transcripts, whereas exon 2 contains both readthrough and non-readthrough signals. Hence, the readthrough fraction was calculated as (depth(exon 2) - depth(exon 1)) / depth(exon 2). (3) Coverage-based ratio for *NPEPPS*: as the terminal exon of *NPEPPS* is partially skipped in the readthrough transcripts, coverage in the unskipped region reflects both transcript types, whereas the skipped segment represents only non-readthrough transcripts. The readthrough proportion was defined as 1 - (depth(skipped) / depth(unskipped)).

Fig. S14: CpG methylation of 37M and 39M from three human individuals.

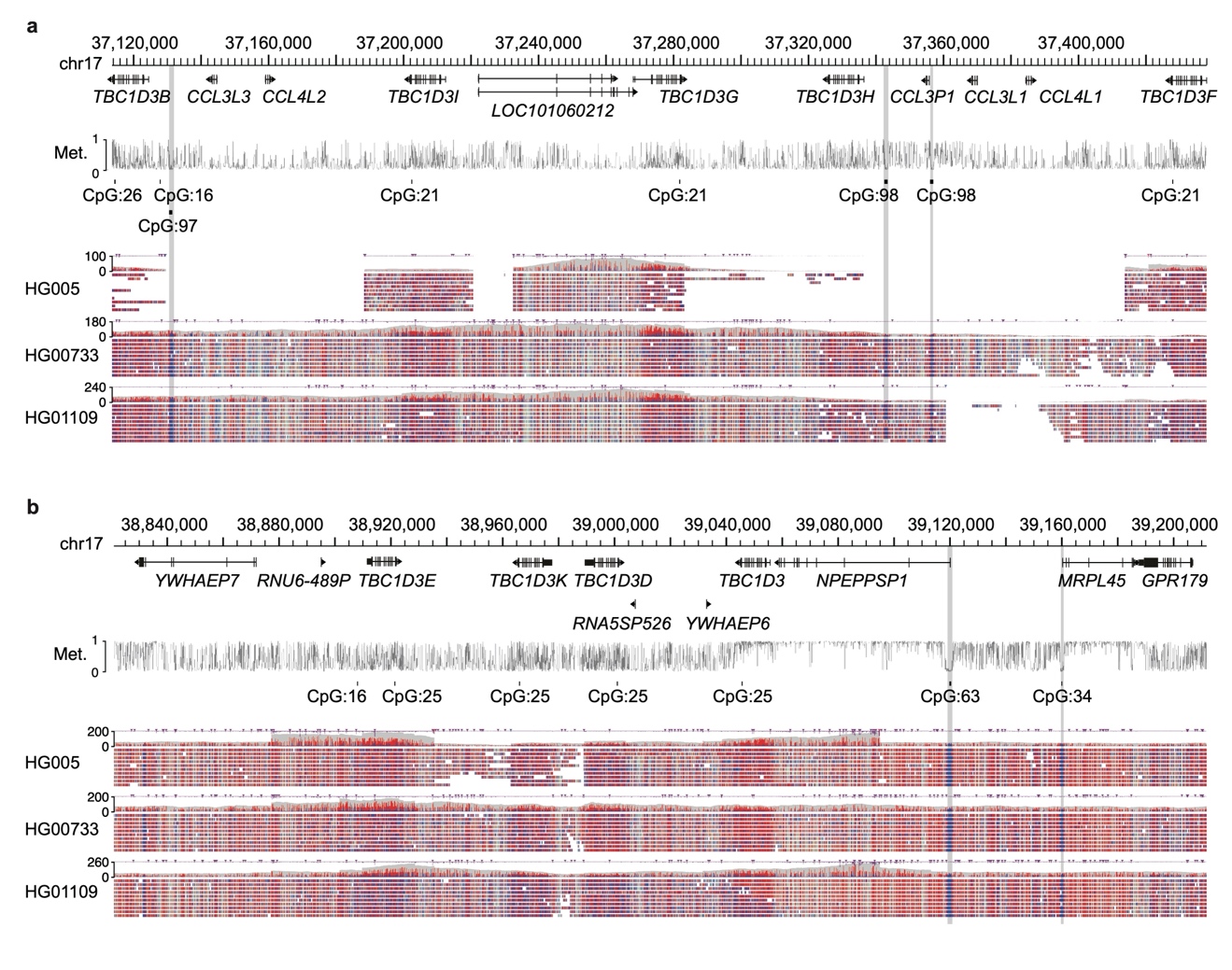

**a**, CpG methylation levels called from HG005, HG00733 and HG01109 aligned nanopore reads indicate that there is no hypomethylated CpG island at the upstream of *TBC1D3* copies at the 37M locus. **b**, CpG methylation levels show the hypomethylated CpG islands (blue) at the upstream of the 39M *NPEPPS*-*TBC1D3* digenic region. The grey line above represents the CpG methylation level of CHM13hTERT cell lines.

Fig. S15: Single-cell analysis of human brain organoid samples shows *TBC1D3*-39M is the predominant paralog for *TBC1D3* expression.

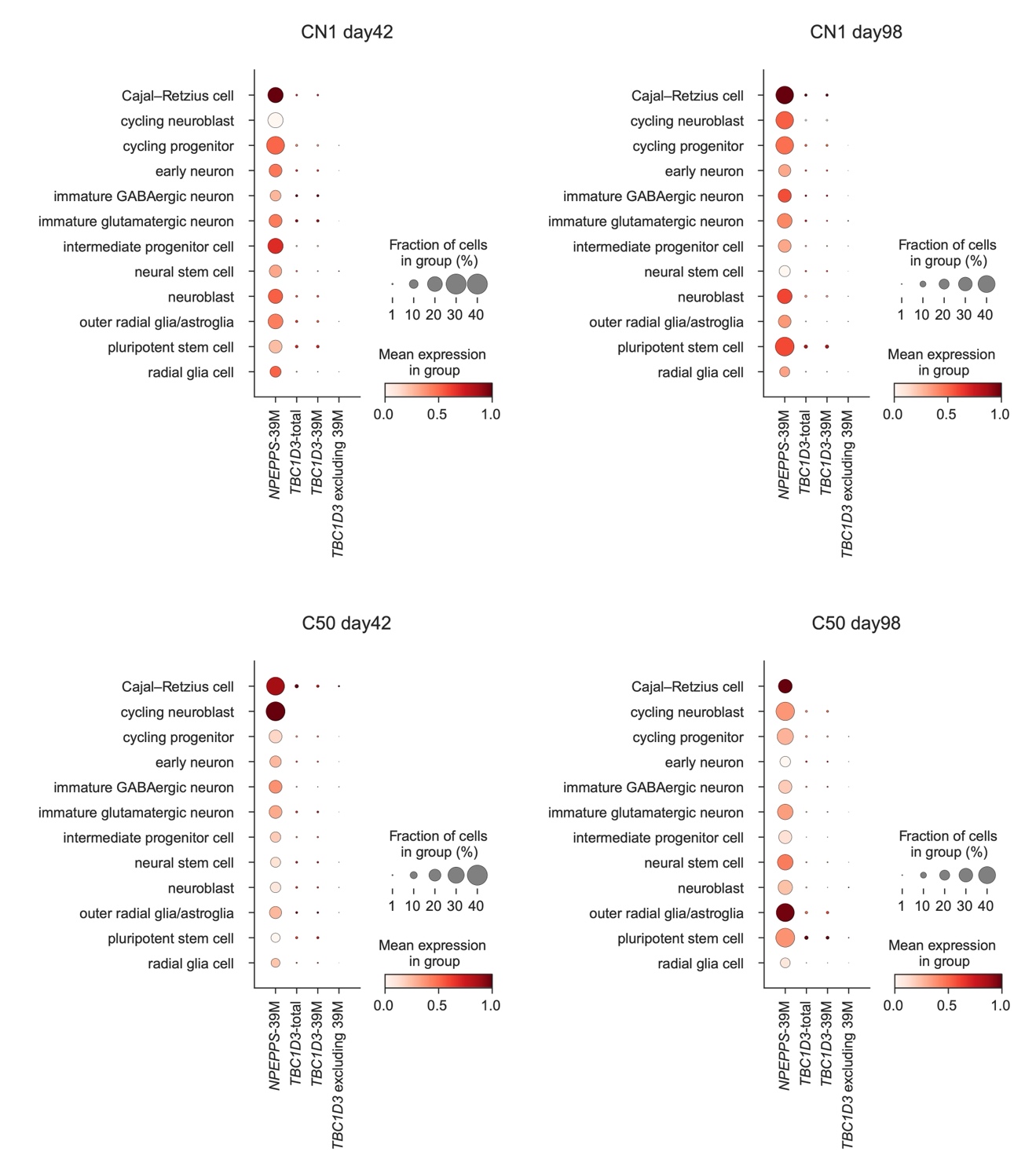

The dot plots show the mean expression of *NPEPPS*-39M, *TBC1D3*-total (sum of all *TBC1D3* paralogs), *TBC1D3*-39M, and *TBC1D3* paralogs excluding *TBC1D3*-39M in a red color scale, while the size of each dot represents the fraction of cells expressing the genes.

Fig. S16: Detailed synteny of 37M, 39M and 48M across primates.

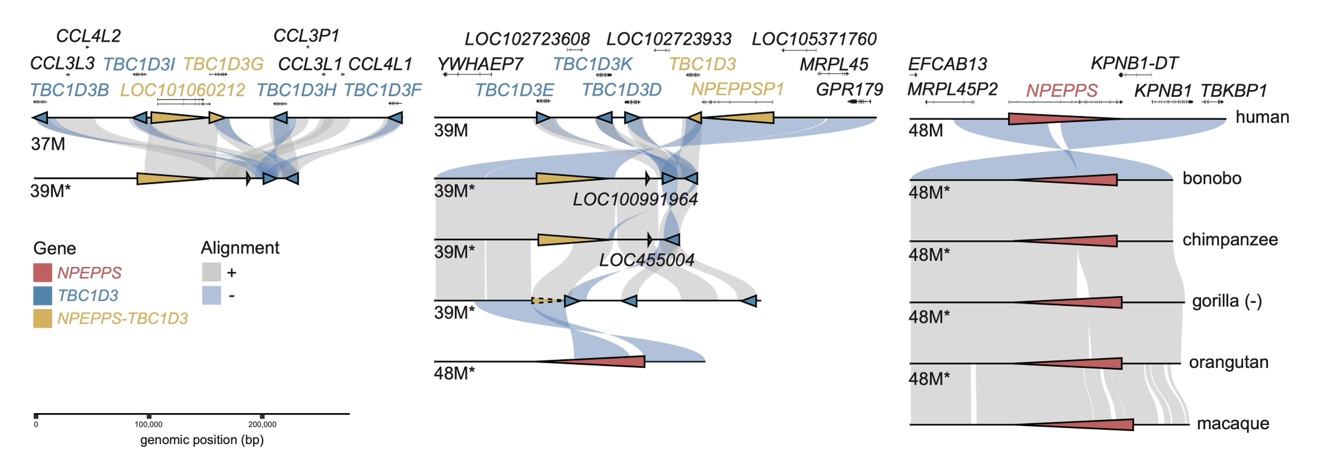

The forward and inverted alignments are shown in grey and light blue, respectively. Red, yellow, blue triangles represent the ancestral *NPEPPS*, duplicated *NPEPPS* and *TBC1D3* copies. The asterisks mark the genomic coordinates of human homologous loci, indicating their corresponding positions in the human genome rather than the native coordinates of non-human primates.
Fig. S17: The unique *NPEPPS*-*TBC1D3* genomic configuration potentially underlies human-specific expression patterns of *TBC1D3*.

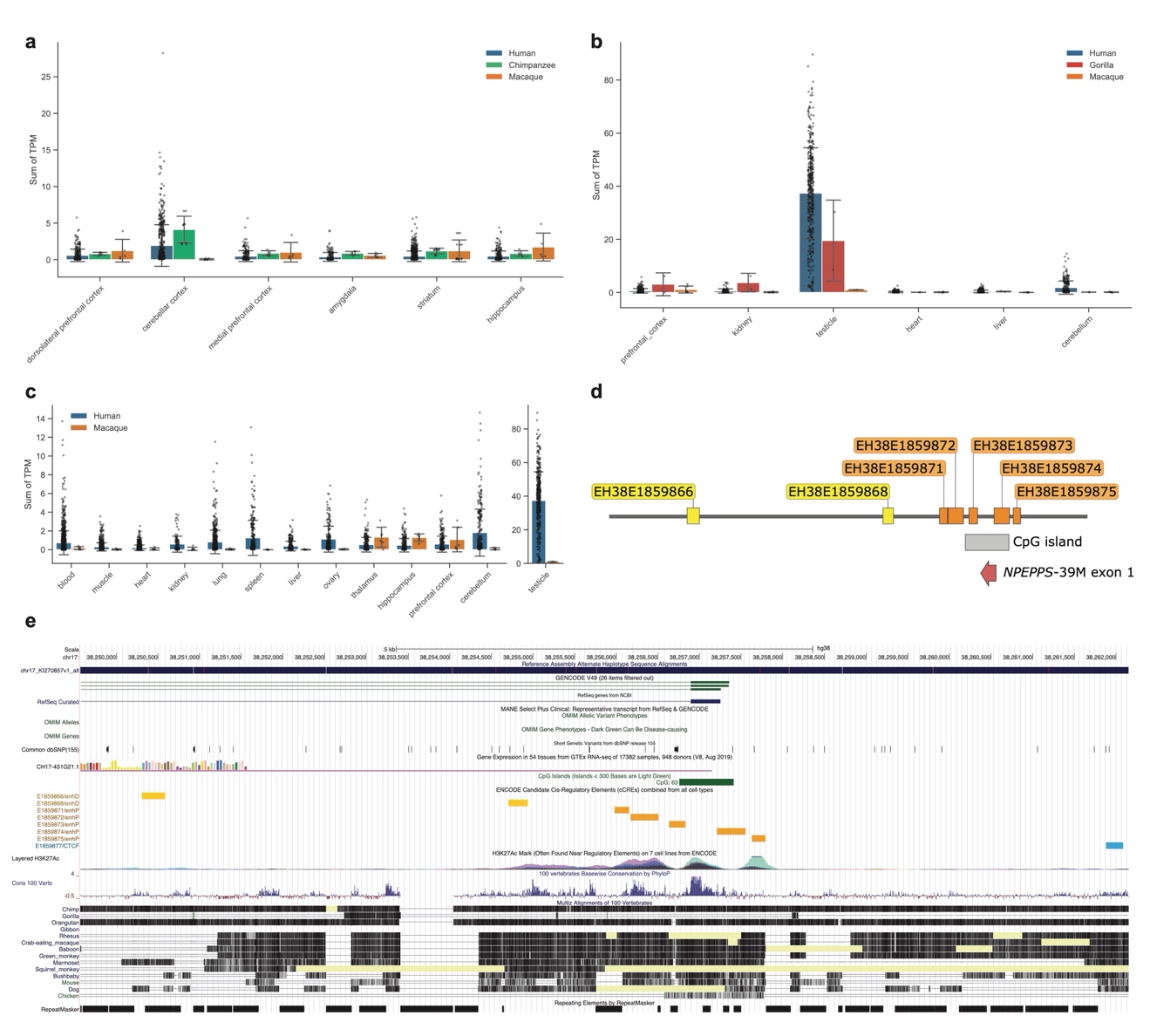

**a**,**b**,**c**, Summed TPM of *TBC1D3* paralogs in humans and NHPs. **d**,**e**, The exon 1 of *NPEPPS*-39M resides within a CpG island and is flanked by candidate *cis*-regulatory elements (cCREs). The cCREs targeting *NPEPPS*-39M presumably regulate the entire readthrough unit, thereby also influencing the expression of *TBC1D3*-39M. TPM matrices for adult human tissues were downloaded from the GTEx Analysis V10 release[^2^](https://sciwheel.com/work/citation?ids=530302&pre=&suf=&sa=0). Raw bulk RNA-seq data for chimpanzee, gorilla, and macaque tissues were obtained from published studies[^3–6^](https://sciwheel.com/work/citation?ids=18441749,4535631,462691,4486888&pre=&pre=&pre=&pre=&suf=&suf=&suf=&suf=&sa=0,0,0,0) (PRJNA1004471, PRJNA143627, PRJNA304995, and PRJNA236446). Raw reads were quality-filtered and trimmed using fastp[^7^](https://sciwheel.com/work/citation?ids=5861897&pre=&suf=&sa=0) (v0.24.1) with default parameters. Clean reads were aligned to their respective reference genomes using STAR[^8^](https://sciwheel.com/work/citation?ids=49324&pre=&suf=&sa=0) (v2.7.11b): NHGRI_mPanTro3-v2.0_pri for chimpanzee (GCF_028858775.2-RS_2024_02), NHGRI_mGorGor1-v2.0_pri for gorilla (GCF_029281585.2-RS_2024_02), and T2T-MFA8v1.1 for macaque (GCF_037993035.2-RS_2025_03)[^9,10^](https://sciwheel.com/work/citation?ids=17734800,17596336&pre=&pre=&suf=&suf=&sa=0,0). Gene annotations were collapsed following the GTEx RNA-seq pipeline (<https://github.com/broadinstitute/gtex-pipeline>), and gene-level quantification was performed using RNA-SeQC[^11^](https://sciwheel.com/work/citation?ids=11844593&pre=&suf=&sa=0) (v2.4.2). For the *TBC1D3* gene family, TPM values across all paralogs were summed.

Fig. S18: (*NPEPPS*-) *TBC1D3* transcript sequences were cloned from SH-SY5Y cells into a pCAGGS expression system.

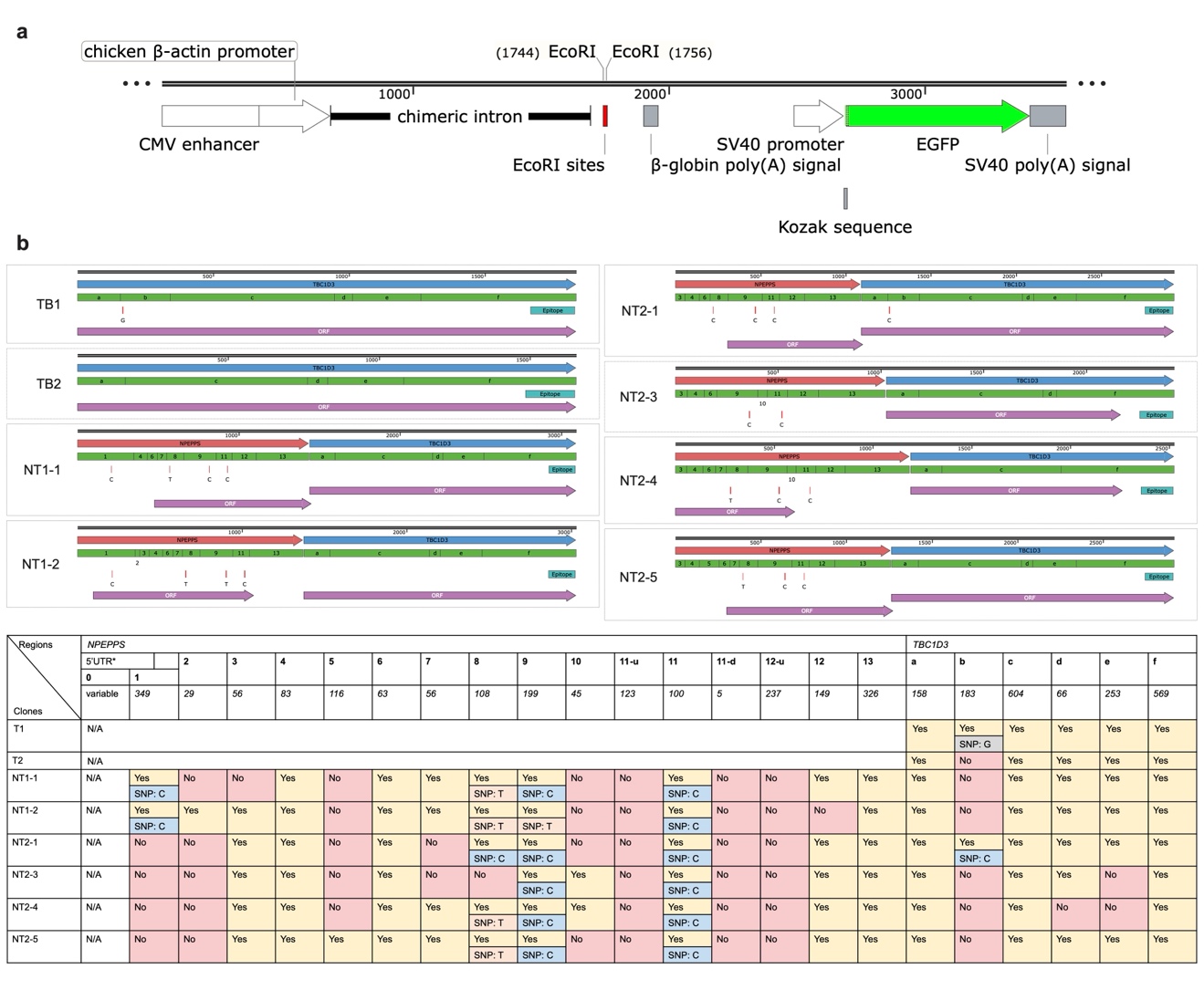

**a**, The transcript sequences were cloned between the EcoRI sites. **b**, TB2 is the same as the ORF of the MANE Select *TBC1D3*. NT1-1 and NT1-2 starts at the predicted TSS of *NPEPPS*-*TBC1D3* while NT2-1, NT2-3, NT2-4, and NT2-5 starts at the annotated start codon of XP_047302714. The NT2-2 was excluded because sequencing showed that the eGFP sequence in the plasmid was disrupted during cloning. 5’ UTR is defined according to: NM_006310.4. N/A: sequence existence is unknown due to primer design. The alternate SNPs are all only observed in one case. It can not be ruled out that they may be PCR artifacts.

Fig. S19: Translational regulations of the *NPEPPS*-*TBC1D3* transcripts.

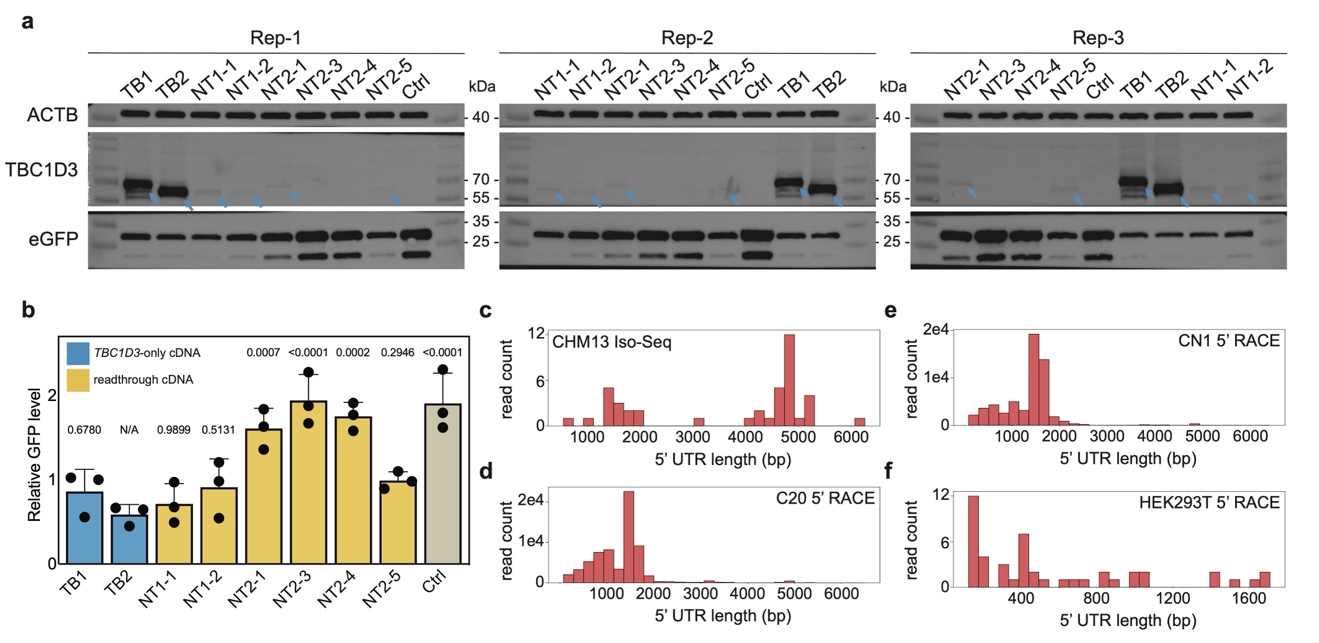

**a**, The readthrough transcripts generate lower TBC1D3 protein level compared to *TBC1D3*-only sequences. **b**, GFP expression is controlled by the SV40 promoter expression cassette in the plasmids, independent from the (*NPEPPS*-) *TBC1D3* expression cassette. ANOVA and Dunnett's multiple comparisons tests were performed for TB2 *vs*. each of the other samples. **c-f**, Distribution of 5’ untranslated region (5’ UTR) lengths for *TBC1D3* readthrough transcripts across four datasets: (c), CHM13 Iso-Seq; (d), C20 5’ RACE; (e), CN1 5’ RACE; (f), HEK293T 5’ RACE. For each dataset, the x-axis represents the length of 5’ UTR (bp), defined as the distance between the readthrough transcription start site and the *TBC1D3* ATG start codon, thereby treating the upstream *NPEPPS* region as part of the 5’ UTR. The y-axis shows the read count supporting each observed 5’ UTR length. Note: Compared to the 5’ RACE results, which may be limited by the extension time, Iso-Seq also detected an additional group of reads retaining a 3.3-kb sequence (T2T-CHM13 chr17: 39068798-39072102) that was annotated as intronic by RefSeq.

Fig. S20: Utilizing currently available tools, no effective IRES was detected in the *NPEPPS* region of the *NPEPPS*-*TBC1D3* readthrough transcripts.

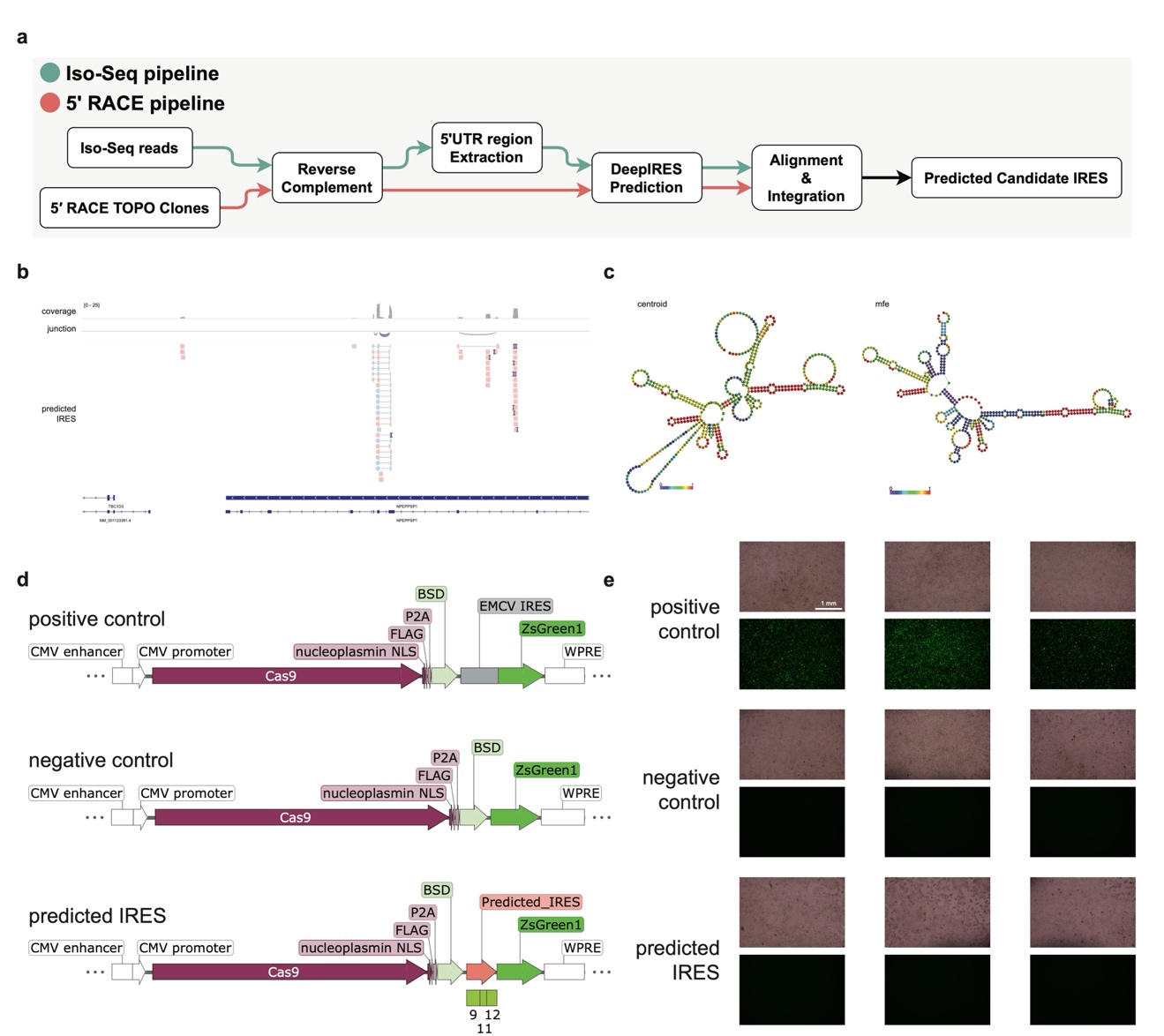

**a,** Overview of the analytical workflow. Iso-Seq reads and 5’ RACE TOPO clones were first processed independently, followed by reverse complementation and extraction of the corresponding 5’ UTR segments. Candidate IRES elements were predicted using DeepIRES[^12^](https://sciwheel.com/work/citation?ids=18448702&pre=&suf=&sa=0), and a final consensus IRES sequence was generated after integrating the Iso-Seq and 5’ RACE information at the last step through sequence alignment. **b**, Genomic visualization of predicted IRES features. IGV tracks display read coverage, splice junctions, and the genomic positions of predicted IRES candidates across the *NPEPPS* region. One consensus sequence derived from the Iso-Seq/5’ RACE co-supporting region was chosen as the candidate predicted IRES, which contains Regions 9, 11, and 12 of *NPEPPS*. **c**, Predicted RNA secondary structures for the candidate predicted IRES. RNAfold results are shown for both the centroid model (left) and the minimum free energy (MFE) structure (right)[^13^](https://sciwheel.com/work/citation?ids=149001&pre=&suf=&sa=0). **d**, a report system was designed and cloned to test the effectiveness of the predicted IRES. **e**, the predicted IRES sequence failed to generate a positive signal (green; as seen in cells transfected with the positive control). The plasmids were transfected into HEK293T cells. Pictures were taken 48 hours post transfection. Three independent transfection replicates were performed for each group.

Fig. S21: The RNA level of the readthrough cDNA is lower than that of the *TBC1D3*-only cDNA using the pCAGGS expression system.

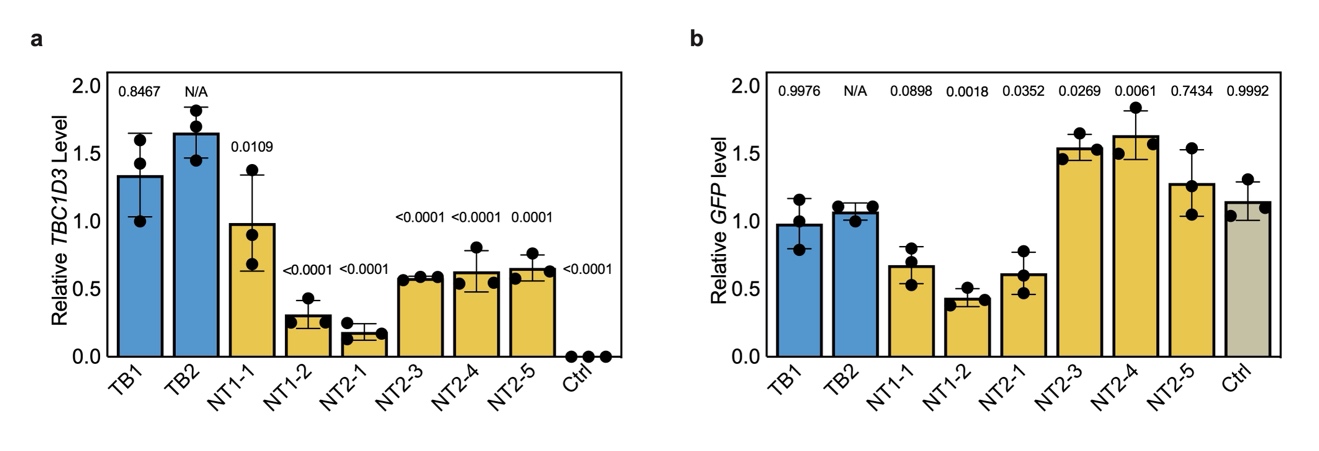

qPCR reactions were performed using the ChamQ Universal SYBR qPCR Master Mix (Vazyme, #Q711) and the following primers: *TBC1D3*-F: 5’-GTGCCATATCCCAGGAGGAC-3’, *TBC1D3*-R: 5’-GCTGTTCGTCCCTAGCTCTG-3’, *GFP*-F: 5’-CCGACAACCACTACCTGAGC-3’, *GFP*-R: 5’-CTTGTACAGCTCGTCCATGC-3’, *HPRT1*-F: 5’-CATTATGCTGAGGATTTGGAAAGG-3’, and *HPRT1*-R: 5’-CTTGAGCACACAGAGGGCTACA-3’. *HPRT1* was used as the housekeeping gene control. qPCR Triplicates were performed. **a**, ANOVA and Šídák's multiple comparisons tests were performed. Adjust p-values for TB2 *vs.* each of the other samples are shown. **b**, ANOVA and Tukey's multiple comparisons tests were performed. Adjust p-values for TB2 *vs.* each of the other samples are shown.

Fig. S22: Ribo-Seq experiments were performed for two representative readthrough transcripts (NT1-2 and NT2-5) and a *TBC1D3*-only transcript.

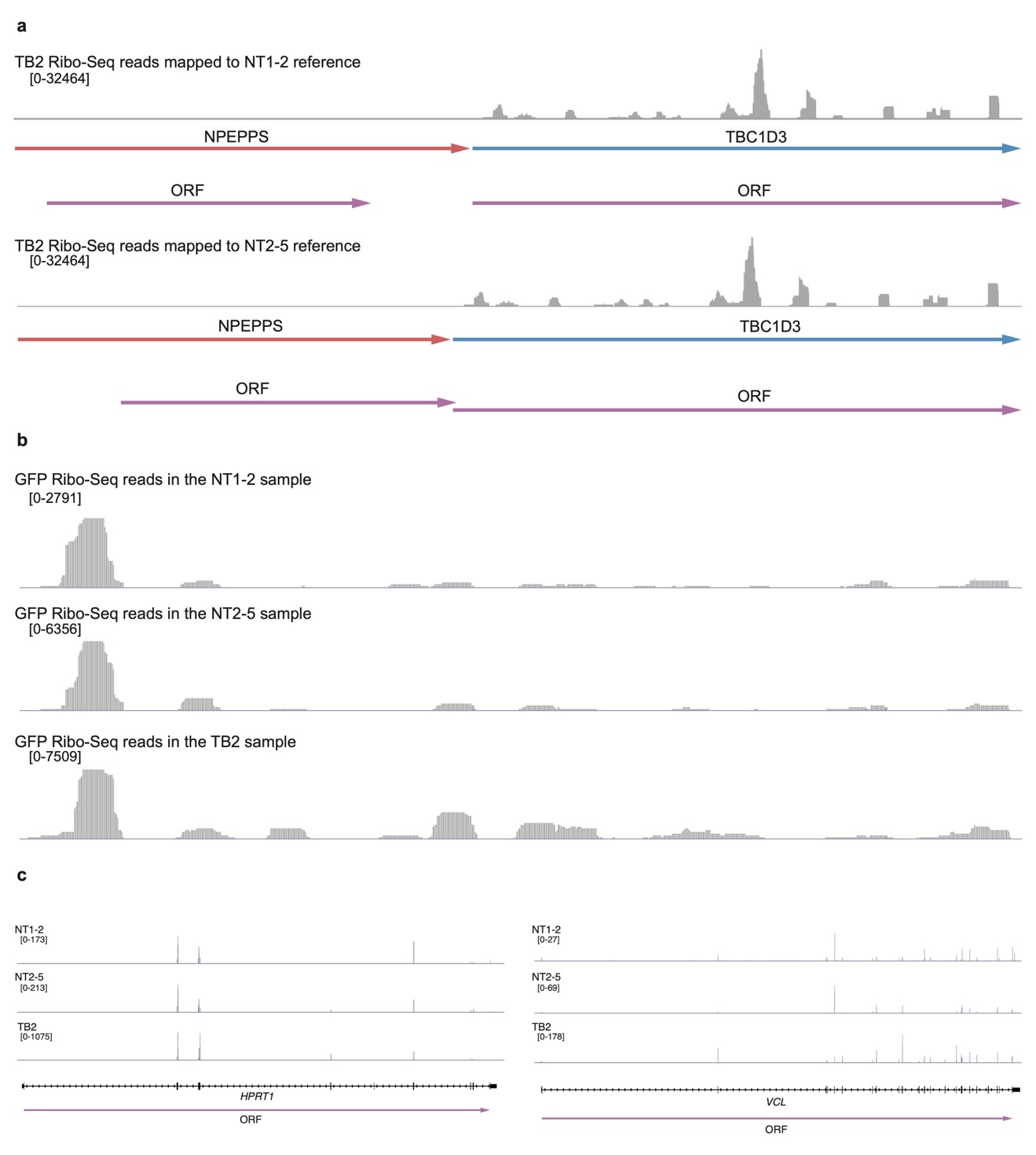

**a**, Mapping the TB2 Ribo-Seq reads to the NT1-2 and NT2-5 references respectively. The endogenous *NPEPPS* expression in HEK293T is neglectable compared to the expression level of the pCAGGS expression system. **b**, The eGFP Ribo-Seq signals of TB2, NT1-2, and NT2-5. **c**, The Ribo-Seq signals of the endogenous *HPRT1* and *VCL* in these 3 samples.

Fig. S23: Genomic, methylation and representative transcript diagrams of JSDTR gene pairs.

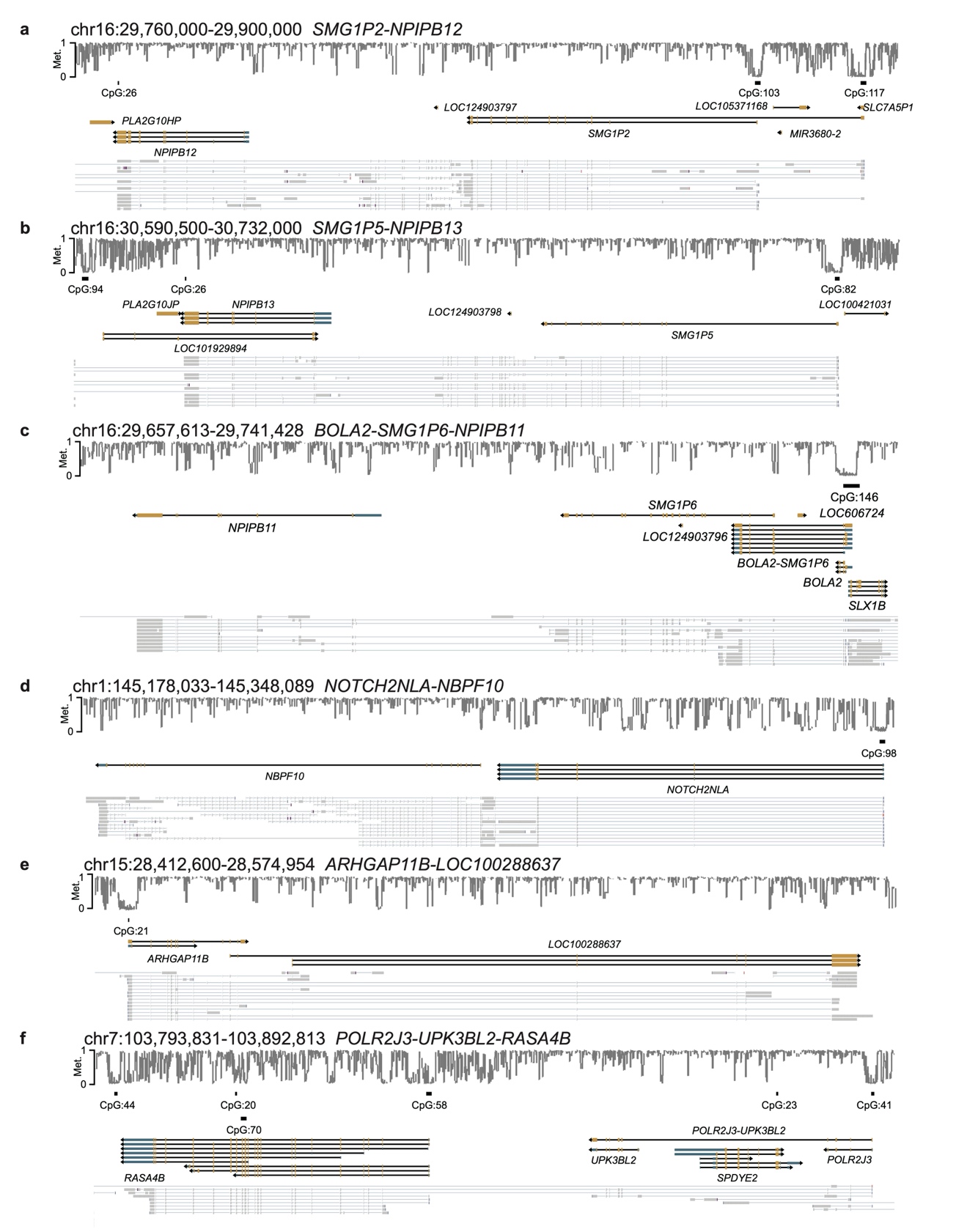

**a-c**, Diagrams of *SMG1P2*-*NPIPB12*, *SMG1P5*-*NPIPB13* and *BOLA2*-*SMG1P6*-*NPIPB11*. **d**, Diagram of *NOTCH2NLA*-*NBPF10*. **e**, Diagram of *ARHGAP11B*-*LOC100288637*. **f**, Diagram of *POLT2J3*-*UPK3BL2-RASA4B* indicates the existence of a JSDTR gene trio.

Fig. S24: *NPEPPS*-39M lacks the poly(A) signal used by *NPEPPS*-48M.

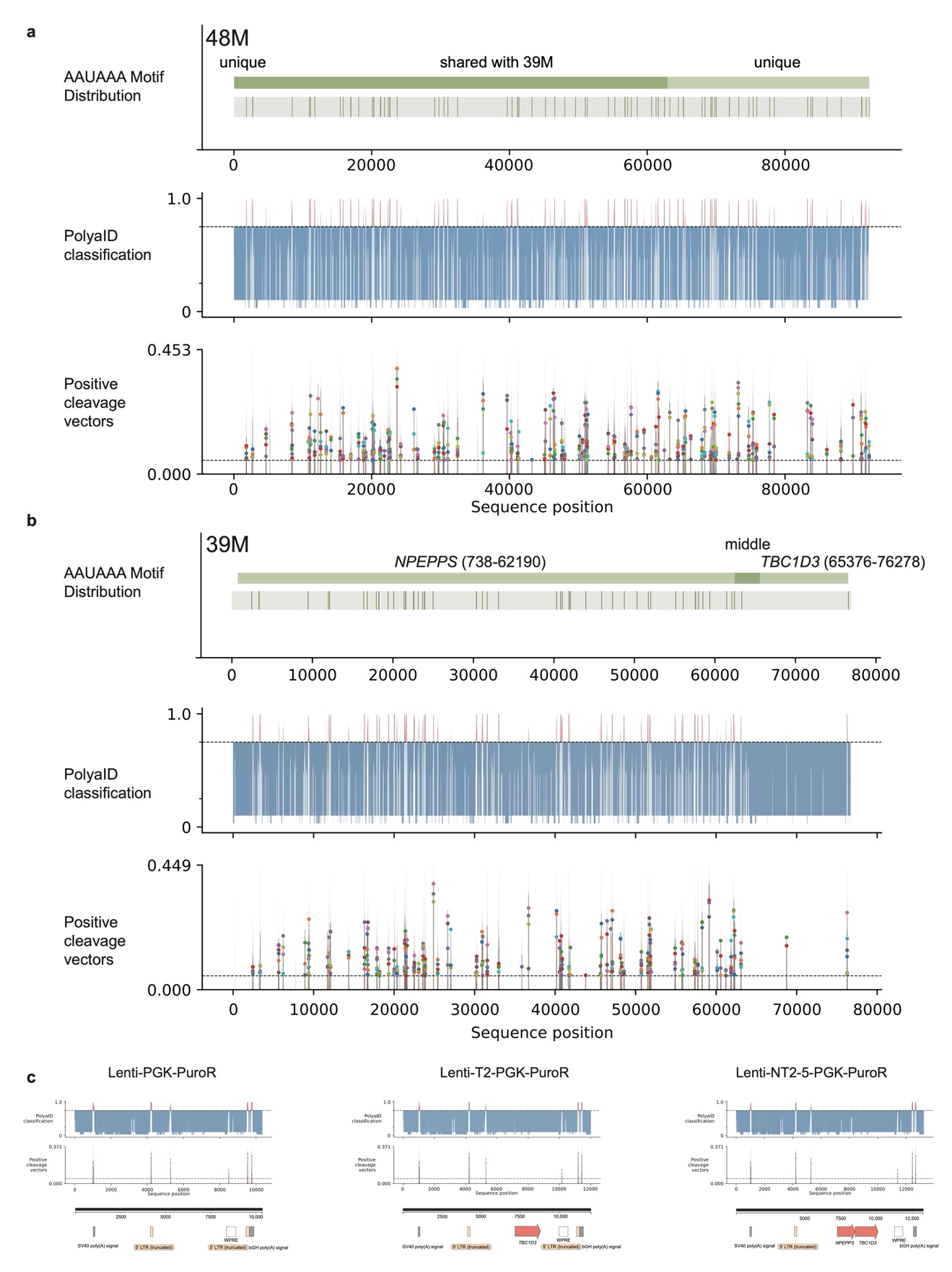

To assess poly(A) signal distribution in the 48M and the 39M loci, we first identified AAUAAA motifs in the sequence and mapped their positions onto the gene structures. We then applied the pretrained PolyaID and PolyaStrength models from zhejilab GitHub (<https://github.com/zhejilab/PolyaModelsHuman>), which predict poly(A) site classification, cleavage profile, and strength score. Input sequences included the *NPEPPS*-48M genomic region (**a**), the *NPEPPS*-*TBC1D3*-39M genomic region (**b**), and three plasmid sequences we designed, which contain commonly used poly(A) signals, as positive controls (**c**). Interestingly, *NPEPPS*-39M contains multiple AAUAAA poly(A) signal motifs, while *TBC1D3*-39M only contains a single motif at the 3' end. Whether this asymmetric distribution also contributes to transcriptional regulation, however, requires further study. Note: In (a), the minor unique region at the 5’ end results more from annotation differences than sequence differences. Note for the PolyaID and PolyaStrength models: Each sequence was extended with 120 "N" bases at both ends, followed by construction of 240-nt sliding windows (1-nt step) and one-hot encoding for model input. PolyaID produced a classification score and a 50-position cleavage vector, with position 25 as the predicted cleavage center. PolyaStrength provided an additional strength score. Cleavage vectors were corrected by subtracting a 0.02 background, zeroing negatives, and normalizing; the center value was recorded as cleavage_center. Windows with classification ≥ 0.75 were considered candidates, and those with cleavage_center ≥ 0.05 were considered high-confidence. Visualization used two panels. The upper panel shows classification scores (blue: <0.75; red: ≥0.75), and the lower displays 50-nt cleavage vectors, with high-confidence vectors (cleavage_center ≥ 0.05) in color with a central marker and low-confidence vectors in faint gray.

Fig. S25: Endogenous TBC1D3 peptides are only occasionally detected.

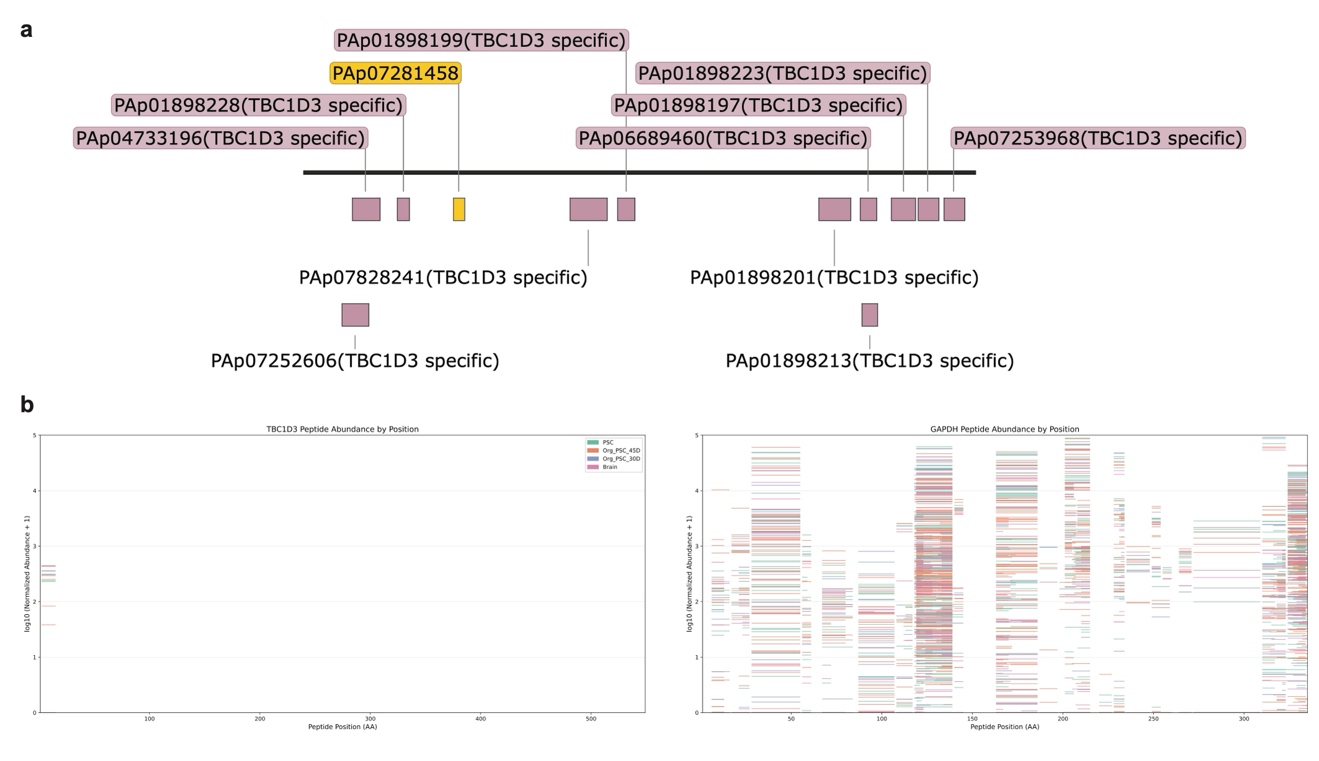

**a**, TBC1D3 peptides recorded in PeptideAtlas are mainly from testis[^14^](https://sciwheel.com/work/citation?ids=1227979&pre=&suf=&sa=0). Here, TBC1D3-specific are the peptides that can only be found in TBC1D3 paralogs but not other proteins. **b**, Peptide abundance mapped along the amino-acid positions of TBC1D3 (left) and GAPDH (right). Each horizontal bar represents the log10-transformed peptide abundance (pseudocounted by +1) detected at the corresponding peptide position. TBC1D3 peptides exhibit sparse and low-abundance detection across all datasets, whereas GAPDH displays dense and relatively uniformly distributed peptide signals.

Fig. S26: Quantification of *NPEPPS-TBC1D3* readthrough and splice-junction evidence in SH-SY5Y cells.

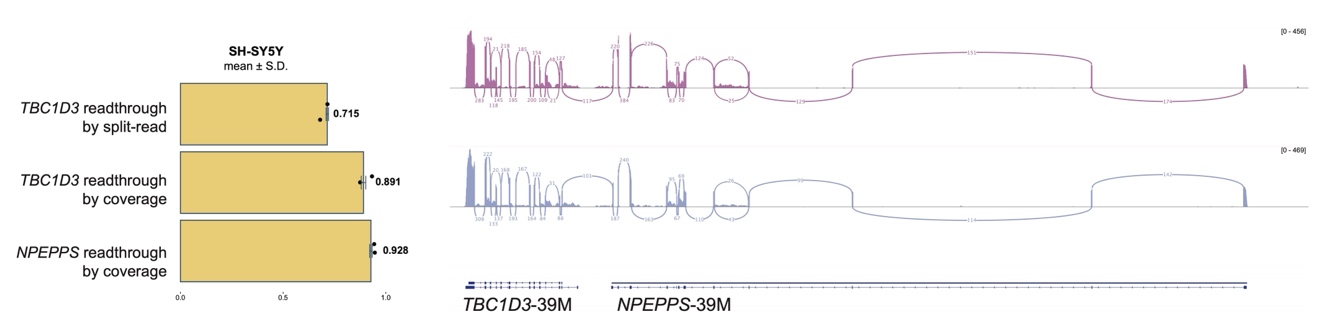

Left: Readthrough ratios of *TBC1D3* and *NPEPPS* in SH-SY5Y cells, quantified using split-read support or junction-spanning coverage. Right: Sashimi plots depicting splice junctions and read coverage across the 39M locus.

Fig. S27: ORFs within the *NPEPPS* region in the readthrough transcripts may share sequences with the annotated Peptidase M1 membrane alanine aminopeptidase domain.

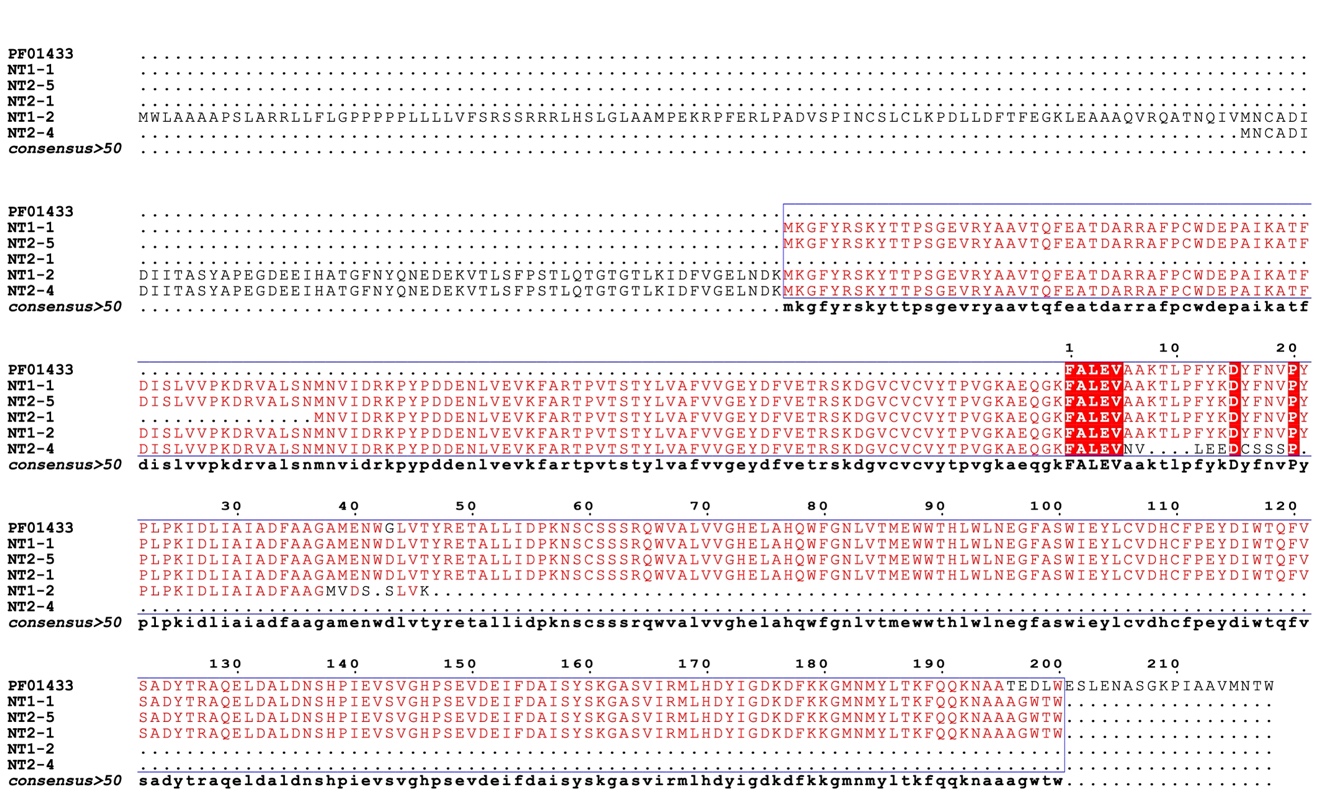

*NPEPPS* ORFs from different cloned *NPEPPS*-*TBC1D3* transcripts share sequences with the annotated Peptidase M1 membrane alanine aminopeptidase domain (PF01433). The comparison was performed with multalin[^15^](https://sciwheel.com/work/citation?ids=1354505&pre=&suf=&sa=0).
